# Plant Cell Wall Enzymatic Hydrolysis: Predicting Yield Dynamics from Autofluorescence and Morphological Temporal Changes

**DOI:** 10.1101/2025.08.11.669614

**Authors:** Solmaz Hossein Khani, Khadidja Ould Amer, Fatemeh Shiasi Ghalemaleki, Maxime Corré, Noah Remy, Anouck Habrant, Berangère Lebas, Jafar Massah, Ali Faraj, Gabriel Paës, Yassin Refahi

**Author notes:** S.HK. and K.OA. contributed equally to this work.

## Abstract

Enzymatic hydrolysis of plant cell walls into fermentable sugars is a critical step in biotechnological conversion, yet efficiency is limited by cell wall recalcitrance. Predicting conversion yields of cell wall-derived sugars during hydrolysis is challenging due to the complex underlying mechanisms and the labor-intensive nature of conventional assays. This study introduces an innovative pipeline that accurately quantifies cell wall autofluorescence intensity and morphological descriptors during enzymatic hydrolysis. The pipeline incorporates a novel adaptive drift compensation strategy which dynamically adjusts to the progression and extent of deconstruction ensuring robust analysis. Applied to time-lapse images of spruce wood enzymatic deconstruction, the pipeline revealed strong negative correlations of conversion yields during hydrolysis with both the dynamics of cell wall autofluorescence intensity and morphological descriptors. Phase-specific analysis uncovered distinct correlation patterns dependent on hydrolysis stage and sugar type. This non-destructive pipeline eliminates the need for extensive sampling and time-consuming chemical assays, establishing plant cell wall autofluorescence and morphological descriptors as accurate predictive real-time biomarkers of dynamics of sugar conversion yields. The findings provide a framework for accelerating the development of optimized biotechnological conversion processes.

## 1 Introduction

Addressing global sustainability challenges such as climate change and depletion of fossil-based resources has intensified the research on using renewable and carbon-neutral resources for energy, chemicals, and materials. Lignocellulosic biomass from plants is the most abundant renewable bioresource with an estimated annual production of around 181.5 billion tons [11]. It can serve as a fundamental component of a suit of solutions, such as wind power and photovoltaic systems, to ensure energy security, and a resilient and sustainable future [9]. The conversion of lignocellulosic biomass such as wood and agricultural residues into bioproducts has an advantage over crop-based feedstock used in first-generation biofuels, such as vegetable oil, and starch, by mitigating food-versus-fuel conflict.

Lignocellulosic biomass is mainly composed of three biopolymers: cellulose (45–55%), hemicellulose (25–35%), and lignin (20–30%) [45]. Cellulose, as the most abundant biopolymer, is a linear polysaccharide composed of *β*-D-glucose monomers linked by *β*(1 → 4) glycosidic bonds. Bundles of aligned *β*(1 → 4)-linked D-glucose chains form cellulose microfibrils [46]. Hemicellulose is an amorphous heterogeneous group of polysaccharides (e.g. xylans, mannans, galactans) composed of various sugar monomers, including pentoses (e.g. arabinose and xylose) and hexoses (e.g. mannose, and galactose) [47]. Lignin is a structurally complex aromatic biopolymer which confers hydrophobicity and mechanical strength to the plant cell walls [57].

Conversion of lignocellulosic biomass into bioproducts involves approaches that can be categorized into biological, chemical, and thermochemical conversions [53]. Biotechnological conversion builds upon biological conversion and offers advantages including mild operating conditions, low toxicity, enhanced selectivity for target product, conversion efficiency and environmental sustainability [10, 56]. These benefits make the biotechnological pathway a promising alternative or complement to thermochemical or chemical conversion methods which use harsher conditions.

The recalcitrance of lignocellulosic biomass poses a significant challenge to its efficient conversion [25, 37] and contributes to the high production cost of bioproducts derived from lignocellulosic biomass. Therefore, to enhance the efficiency and economical viability of transformation of lignocellulosic biomass, the recalcitrance should be addressed. Key factors contributing to recalcitrance include the chemical composition and structural characteristics of lignocellulosic biomass. The interactions between hemicelluloses, cellulose, and lignin limits the enzyme accessibility and enzymatic hydrolysis efficiency. Hemicelluloses create an enveloping matrix around cellulose microfibrils forming a physical shield that restricts enzyme access. Structural and physical properties of cellulose, in particular crystallinity and degree of polymerization, play a significant role in biomass recalcitrance [28]. High cellulose crystallinity and high cellulose degree of polymerization negatively impact the enzymatic hydrolysis efficiency [23]. Lignin related characteristics of lignocellulosic biomass are widely recognized as major contributors of recalcitrance [57]. These characteristics include lignin content, lignin composition, as relative proportions of S, G, and H units in the total lignin subunits, molecular weights (such as weight-average molecular weight, and polydispersity index) and phenolic hydroxyl groups [57]. In addition to making a natural barrier to cellulose, lignin also contributes to creating non-specific interactions with enzymes, reducing their efficiency [17, 57, 58]. In addition, structural features of lignocellulosic biomass such as pore size and cellulose accessibility, also referred to as accessible surface area of cellulose, play a crucial role in recalcitrance [24, 38, 39].

Biotechnological conversion of lignocellulosic biomass involves three main steps: pretreatment, enzymatic hydrolysis, and fermentation [11]. Pretreatment disrupts the complex structure of the biomass and reduces the biomass recalcitrance. Pretreatment is followed by enzymatic hydrolysis, a key bottleneck limiting the efficiency of lignocellulosic biomass biorefineries, to generate simple fermentable sugars [21]. The sugars are then converted into bioproducts, like bioethanol, via fermentation by yeast, bacteria, and fungi, through separate or integrated process strategies [36]. Enzymatic saccharification can be considered the most critical step [1, 21, 50]. Effective enzymatic hydrolysis is not only essential for achieving optimal yields but also has a decisive role in determining the economic viability of producing bioethanol from lignocellulose [6]. However, dynamics of conversion yields of cell wall-derived sugars remain elusive to predict, primarily due to cell walls complex and heterogeneous nature. In addition, the yield conversion measurements remains time-consuming and laborious, often relying on traditional two-stage sulfuric acid hydrolysis followed by HPLC or enzyme-linked spectrophotometric assays, such as glucose oxidase or DNS-based methods, which involve extensive manual handling [52]. The unpredictability combined with low throughput not only impedes rapid screening of biomass feedstocks and process conditions in research contexts but also constrains industrial efforts to optimize enzyme formulations and reaction parameters in a cost-effective manner.

Study of plant cell wall structure that govern its susceptibility to enzymatic deconstruction increasingly relies on advanced imaging and spectroscopic techniques such as atomic force microscopy (AFM), AFM-based approaches such as nano-FTIR (Fourier transform infrared spectroscopy) [5, 31] and confocal Raman microscopy[20, 60], which have advanced our understanding of the chemical composition, structural heterogeneity, and hydrolysis efficiency of plant cell walls. Solid-state NMR techniques have contributed to understanding of the molecular architecture of plant cell walls and elucidating specific interactions such as those between cell wall components [19, 30, 51, 59]. While most investigations are at nanoscale, microscale studies have also advanced our understanding of cell wall enzymatic hydrolysis. Fluorescent probes, immunolabeling, and cellulose staining have been employed for imaging of enzymatic hydrolysis, offering visual insights into cell wall polymer deconstruction and its dynamics [32]. Furthermore, recent studies have also demonstrated the relevance and importance of microscale quantitative investigation of cell walls enzymatic hydrolysis [26, 49]. Despite progress, cell and tissue scale investigation of morphological parameters of plant cell wall during enzymatic deconstruction remains challenging and quantitatively underexplored. This is because acquiring time-lapse image datasets is notoriously demanding, with only a few currently available in public repositories [27]. More importantly, accurate quantitative analysis of time-lapse images is particularly challenging, with sample drift being one of the common challenges to be addressed. A widely used strategy for drift compensation relies on fiducial markers, such as gold nanoparticles or fluorescent beads whose positions are tracked to quantify three-dimensional drift [4, 34]. Alternatively, marker-free drift correction methods have been developed, offering the advantage of eliminating the need for specialized sample preparation or modifications to the imaging system [13]. Existing drift compensation approaches include [26, 49]: i) global approach which registers images to a single reference image but fails under extensive hydrolysis; ii) local approach which registers consecutive images which is suitable for handling substantial changes between images under high deconstruction conditions, but accumulates error through repeated transformation compositions; and iii) fixed-size clustering approach which divides time-lapse images into clusters of predefined sizes. This approach offers to balance prior approaches and uses a global approach, used for intra-cluster images, and local approach across the clusters. However, this clustering strategy requires manual tuning and is suboptimal under natural variable deconstruction rates, often resulting in excessive registrations and reduced accuracy.

This study introduces AIMTrack (Adaptive autofluorescence Intensity and Morphology Tracking), a novel computational pipeline devised to quantify dynamics of autofluorescence intensity and morphological changes of cell walls during enzymatic deconstruction. AIMTrack employs an adaptive clustering strategy to compensate for sample drift and deformation where the cluster sizes are adapted to deconstruction.

Time-lapse images of pretreated spruce wood are then collected. The pretreatment is preformed using sodium chlorite to achieve partial delignification [22]. Spruce was chosen due to the impact of climate change on European forests experiencing widespread spruce dieback [43]. AIMTrack is applied to the collected time-lapse images of pretreated spruce wood, and dynamics of enzyme-driven changes in cell wall autofluorescence and morphological descriptors are quantified. Finally, the relationship between quantified dynamics and cell wall monosaccharide conversion yield dynamics is investigated.

## 2 Results

### 2.1 Dynamics of enzymatic hydrolysis in raw and chlorite-treated spruce wood samples

Chemical compositions of the raw and sodium chlorite–pretreated spruce wood samples were determined to assess the extent of chemical changes of the pretreatment process (Fig. S1). The raw sample was mainly composed of glucose (46.35%), lignin (30.42%), mannose (11.76%), xylose (5.45%), galactose (2.63%) with only minor amounts of other sugars including arabinose, galacturonic acid, glucuronic acid, rhamnose, and fucose. In contrast, the compositional analysis of the chlorite-pretreated sample revealed a remarkable increase in the proportion of glucose (58.89%). Mannose, xylose, and galactose content also shifted to 15.01%, 6.5%, 1.44% respectively. Lignin content dropped to 11.26% following pretreatment, in line with expected compositional shift, while contents of other sugars present in small amounts changed slightly.

To characterize the kinetics of enzymatic hydrolysis, conversion yields of the sugars identified in the compositional analysis were measured during 24 h for both pretreated and raw samples under three experimental conditions: a control without enzymes, and two enzyme loadings of 15 and 30 FPU/g biomass. Significant conversion yields were observed in pretreated samples under both enzyme loadings (Fig. S2). In contrast, only minor saccharification was observed in the raw samples at both enzyme loadings (Fig. S3). In both pretreated and raw samples under control conditions, sugar conversion yields were negligible.

More specifically, pretreated samples subjected to enzymatic hydrolysis at 30 FPU/g biomass consistently exhibited the highest conversion yields at every time point. Conversion yield of glucose showed fast increase during the first 4 h and started to plateau after 8 h. Similarly, xylose exhibited an initial high rate of enzymatic hydrolysis, with a slower, but steady conversion rate over the subsequent 20 h. In contrast, mannose and galactose conversion yields maintained consistently steady dynamics throughout the hydrolysis process over 24 h. Analysis of grouped hemicellulosic sugars conversion yields revealed a consistent increase over time with a moderately accelerated conversion rate during the first 4 h. However, this early-stage trend was less pronounced than that seen for glucose over the same period. Pretreated samples hydrolyzed at 15 FPU/g biomass exhibited similar trends of conversion yields dynamics, but with lower conversion yields compared to pretreated samples hydrolyzed at 30 FPU/g biomass. Altogether, these results highlight an enzyme activity-dependent effect: higher enzyme loading (30 FPU/g biomass) leads to faster and more complete sugar conversion, while lower enzyme loading (15 FPU/g biomass) achieves a significant, but comparatively lower yields. Furthermore, while glucose conversion reaches a plateau after approximately 8 h, hemicellulosic sugars conversion continues to progress after that time frame.

### 2.2 Adaptive pipeline to track cell wall autofluorescence intensity and morphology during hydrolysis

To assess the impact of enzymatic hydrolysis on plant cell wall, confocal fluorescence time-lapse 3D images of pretreated spruce wood samples were collected over the course of hydrolysis, with two enzyme loadings of 15 and 30 FPU/g biomass utilizing the inherent autofluorescence of cell wall [16]. Additionally, control time-lapse images were acquired under the same conditions but without enzymes. Time-lapse imaging and the subsequent analysis were limited to pretreated samples, as raw samples exhibited only minor saccharification. The images were acquired with a one hour interval over 24 h. Hereafter, the control time-lapse images are referred to as *control dataset*, and those collected with two enzyme loadings of 15 and 30 FPU/g biomass are collectively referred to as *hydrolysis dataset*. The first image in a time-lapse series is designated as pre-hydrolysis image and the following images are referred to as hydrolysis-phase images. For consistency, the term hydrolysis-phase images is also used for the post-initial time-lapse images in the control dataset, even though no enzymatic hydrolysis occurs.

The hydrolysis dataset revealed highly deconstructed cell walls exhibiting material loss, structural deformations, a progressive decline in autofluorescence intensity and gradual sample drift during timelapse imaging (Fig. 1). Consequently, tracking cell walls undergoing deconstruction from time-lapse images required efficient drift and deformation compensation to ensure accurate reliable quantification.

**Fig. 1:**
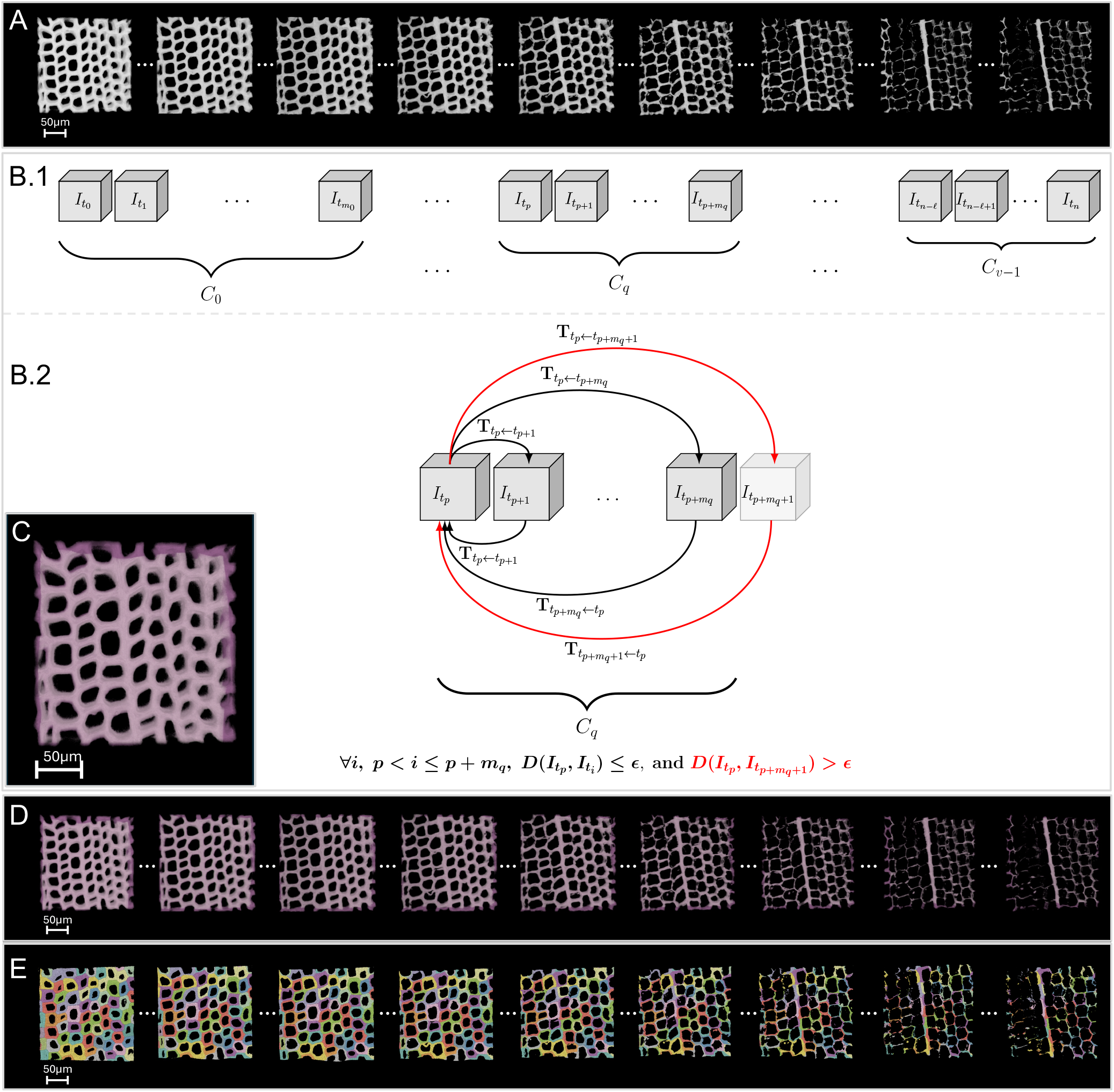
Tracking of cell wall during enzymatic hydrolysis. (A) Time-lapse images of cell wall hydrolysis. (B) AIMTrack’s adaptive clustering framework: (B.1) 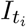 is the *i*-th 3D image collected at time *t*_*i*_, 0 ≤ *i* ≤ *n*. AIMTrack’s variable cluster size strategy employs larger clusters during periods of slower cell wall modification, and adaptively reduces cluster size during periods of faster structural changes. (B.2) AIMTrack ensures that deformation magnitude, *D*(.,.), within each cluster does not exceed a predefined threshold, *ϵ*. The last image of a cluster is the final image in the contiguous sequence for which the deformation does not exceed *ϵ*; the image immediately following exceeds this threshold. The transformations are computed in forward and backward directions within each cluster which are used to evaluate structural changes induced by enzymatic hydrolysis. Composition of intra-cluster transformations registers hydrolysis-phase images to the pre-hydrolysis image, effectively compensating for drift and deformation. (C) Pre-hydrolysis cell walls consistently captured throughout the time-lapse are rendered with increased brightness. (D) Corresponding tracking of these cell walls during hydrolysis, with enhanced brightness indicating their position at each time point. (E) Identification of individual cells in the time-lapse images.

The existing drift compensation methods demonstrated limitations and reduced accuracy. The global approach which involves registering the hydrolysis-phase images directly with pre-hydrolysis, although initially effective, failed as deconstruction advanced and hydrolysis-induced changes accumulated. Local approach which involves indirect registration of hydrolysis-phase images and pre-hydrolysis image by registering pair of consecutive images and composing the resulting transformations, became inefficient by accumulating errors from repeated registration and compositions. The clustering approach is a compromise between the global and local approaches by limiting the registration to clusters, thus reducing registration time-frame compared to global approach, and reducing the required number of transformation composition compared to local strategy [26]. However, clustering strategy with fixed-size clusters has some limitations. The optimal cluster size selection relies on iterative visual assessment of registration quality which can be a laborious and subjective process. Moreover, the optimal cluster size has to minimize the registration and transformation composition frequency, therefore needs to be as large as possible without comprising the intra-cluster registration precision. However, considerable temporal variability is observed in hydrolysis rates and associated changes in cell walls [26]. Significant cell wall modification induced by enzymatic hydrolysis within short time intervals, necessitates a smaller cluster size to achieve precise registration. Using a fixed cluster size implies the consistent use of this smaller cluster size throughout the entire sequence of time-lapse images, even during intervals lacking significant cell wall modifications. This results in an increased number of registration and transformation compositions, and reduces the quality of indirect registration.

To address these limitations, an adaptive clustering method named AIMTrack (Adaptive autofluorescence Intensity and Morphology Tracking) was developed to track cell wall autofluorescence intensity and morphodynamics during enzymatic hydrolysis. AIMTrack includes a hybrid drift and deformation compensation strategy by using a global strategy within clusters and a local strategy between clusters (Fig. 1.B and Fig. S4.A) and introduces adaptivity by dynamically adjusting cluster sizes according to the extent of deconstruction with larger clusters for slow deconstruction stages and smaller clusters for stages involving more significant deconstruction. This adaptive clustering approach builds upon the divide-and-conquer principle [12] by adopting a dynamically modulated partitioning strategy. More precisely, AIMTrack initiates clustering from the pre-hydrolysis image and assesses deformation between the pre-hydrolysis image and subsequent hydrolysis-phase images via registration (Fig. S4.A). The cluster progressively expands with images exhibiting only gradual changes in average deformation. A pronounced increase in average deformation indicates the limit of effective registration and consequently marks the end of cluster expansion. AIMTrack initiates a new cluster by including the final image of the previous cluster and image associated with the rapid average deformation increase, and the same process used for determining the preceding cluster repeats to determine the current cluster’s right-most image. This process repeats until reaching the last image of the time-lapse dataset. The resulting transformations are then used to compensate for sample drift and deformation enabling identification of the consistently imaged region, defined as the intersection of resampled hydrolysis-phase images in frame of the pre-hydrolysis image. The cell walls within this region, called consistently imaged cell walls (Fig. 1.C), are then propagated across the hydrolysis-phase images using the transformations enabling tracking of cell walls and their autofluorescence intensity over time (Fig. 1.D and Fig. S4.B). Furthermore, this propagation strategy is also used to identify the individual cell walls during hydrolysis using their initial states in pre-hydrolysis image (Fig. 1.E and Fig. S4.C).

To assess the performance of the AIMTrack adaptive clustering, its output was compared with expert determined clusters based on visual inspection. This comparison revealed strong agreement with approximately 95% of the images clustered by AIMTrack matching the expert annotations (Table 1). When compared with fixed-size clustering, AIMTrack generated larger average cluster sizes, particularly for the hydrolysis datasets (Fig. 2). At 30 and 15 FPU/ g biomass, AIMTrack provided larger clusters (5.23 and 13, respectively), compared to the fixed-size cluster method (2.66 and 8.00, respectively). For the control dataset, AIMTrack produced an average cluster size of 21.0, closely matching the fixed-size strategy (20.0). Larger cluster sizes reduce the frequency of registrations and transformation compositions, therefore minimize cumulative error and improve overall accuracy [26]. The benchmarking metrics, including agreement with expert annotations, manual intervention requirements, and error impact, are summarized in Table 1. Collectively, these results demonstrated the ability of AIMTrack in autonomously determining the clusters, eliminating the need for manual intervention, and enhancing precision. Furthermore, the larger clusters generated by AIMTrack, and the correspondingly reduced number of clusters, results in enhanced accuracy of segmentations of hydrolysis-phase images as previously demonstrated [26].

**Table 1:** Quantitative benchmarking of AIMTrack adaptive clustering. Metrics include agreement with expert annotations, average cluster sizes, requirement for manual intervention, and impact on cumulative registration error against fixed-size clustering. Cluster size values correspond to those shown in Fig. 2 but are provided here for precision and to facilitate direct comparison with additional performance indicators.

| **Metric** | **AIMTrack** | **Fixed-size Clustering** | **Improvement** |
| --- | --- | --- | --- |
| Agreement with expert annotations (%) | 95 | – | High accuracy |
| Average cluster size (30 FPU g<sup>-1</sup> biomass) | 5.23 | 2.66 | ×1.97 |
| Average cluster size (15 FPU g<sup>-1</sup> biomass) | 13.00 | 8.00 | ×1.63 |
| Average cluster size (control dataset) | 21.00 | 20.00 | ×1.05 |
| Manual intervention required | No | Yes | Eliminated |
| Impact on cumulative error | Reduced | Higher | Improved precision |

**Fig. 2:**
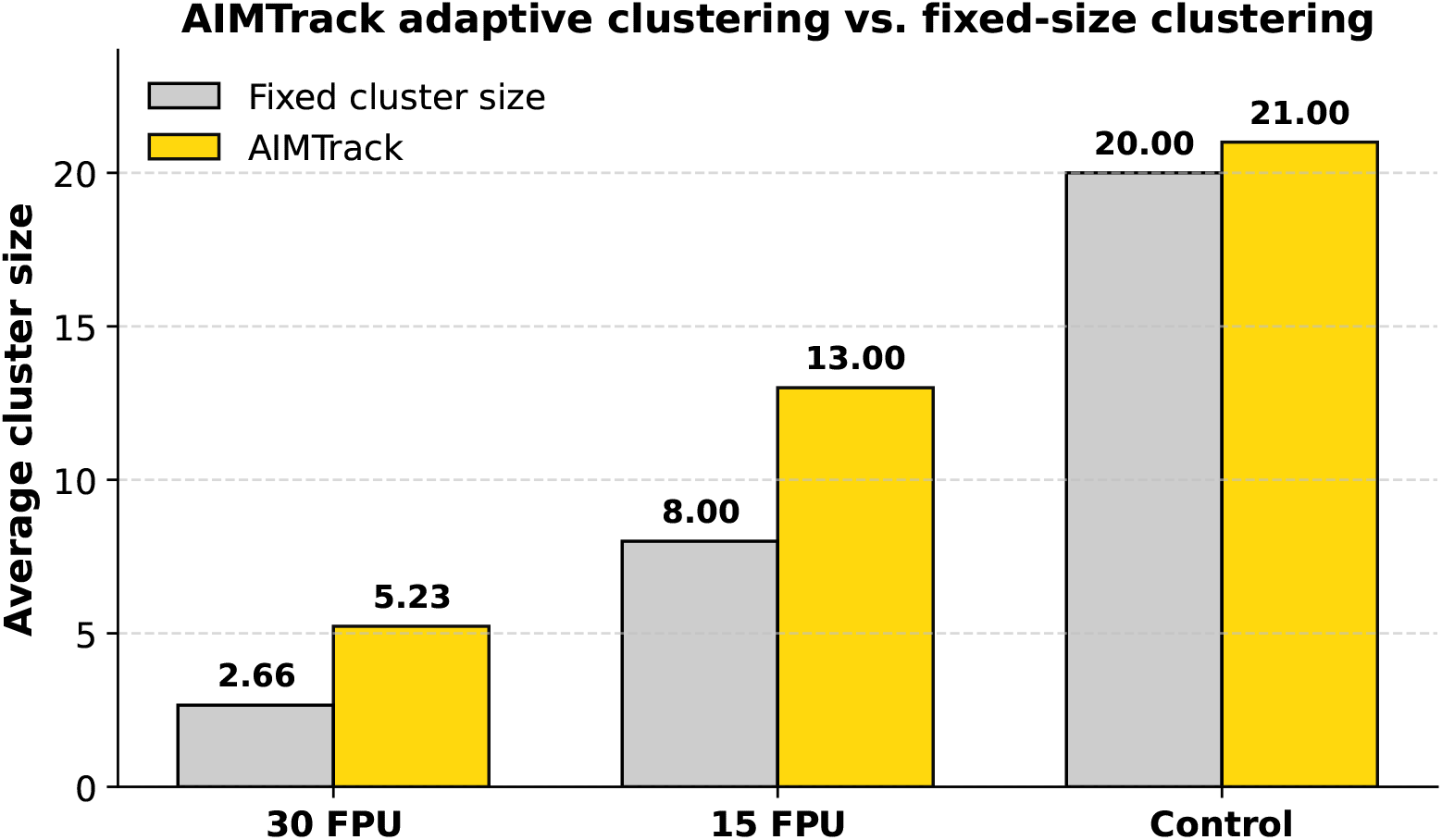
Comparison of average cluster sizes produced by fixed-size clustering and AIMTrack under different enzymatic conditions. In the presence of enzymes at enzyme loadings of 30 FPU/g biomass and 15 FPU/g biomass, AIMTrack provides larger clusters than fixed-size clustering strategy. Both strategies generate comparable cluster sizes for control time-lapse datasets. By producing larger clusters where appropriate, AIMTrack reduces the number of transformation compositions, therefore improves accuracy.

### 2.3 Enzyme-driven cell wall autofluorescence intensity dynamics are monotonic and dependent on enzyme loading

To investigate the cell wall autofluorescence intensity evolution during enzymatic hydrolysis, normalized average autofluorescence intensity dynamics, hereafter referred to as AFI dynamics, were first computed directly from the unprocessed control and hydrolysis datasets (Fig. 3.A). The computed AFI dynamics exhibited gradual decreases, with control datasets showing the lowest drop compared to the hydrolysis dataset. The collected time-lapse images were then processed using AIMTrack to correct the sample drift and compute the AFI dynamics of the consistently imaged region (Fig. 3.B). The AFI dynamics showed comparable overall trends between the unprocessed and drift-corrected image datasets. In both cases, AFI dynamics of the hydrolysis dataset showed larger reductions than the control. However, the drift-corrected datasets undergoing enzymatic hydrolysis displayed a more pronounced AFI dynamics decline compared to the unprocessed hydrolysis datasets without drift correction (Fig. 3.A, B). The drift correction ensured a more precise extracted AFI dynamics by focusing on the same spatial region of the sample which is common to all images, the consistently imaged region, and removing the inaccuracy induced by positional inconsistency. Following the identification of the consistently imaged region in the pre-hydrolysis images, the consistently imaged cell walls within this region were isolated by removing background noise. AFI dynamics quantification was restricted to these consistently imaged cell walls, meaning that only time-dependent changes in autofluorescence intensity within this region were tracked (Fig. 3.C). Consistent with the previously observed trends, the computed AFI dynamics displayed a gradual decline over time, with control datasets exhibiting the smallest reductions and the hydrolysis dataset demonstrating more significant decreases. However, the decrease in autofluorescence intensity became markedly more pronounced under different experimental conditions. Specifically, distinctions in AFI dynamics between control and hydrolysis datasets became clearer, with higher enzyme loadings exhibiting a distinctly greater decline. This exclusive focus on cell wall voxels (3D pixels) enabled to effectively isolate the cell wall-specific signal by discarding the background and avoiding underestimation or misinterpretation of AFI dynamics.

**Fig. 3:**
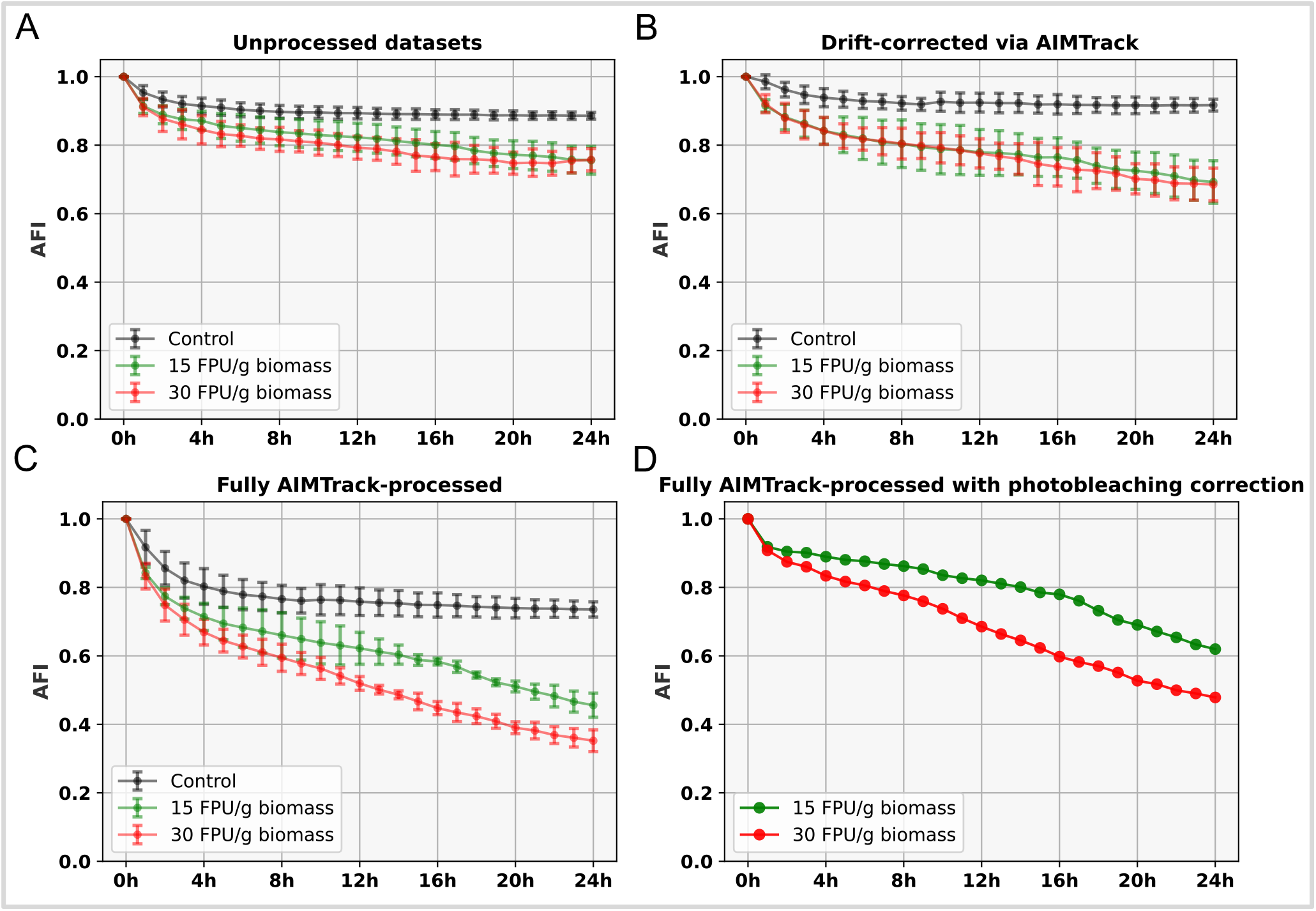
Autofluorescence intensity dynamics (AFI). (A) Normalized AFI dynamics extracted from original (unprocessed) datasets. (B) Normalized AFI dynamics after drift correction using AIMTrack pipeline. (C) Normalized AFI dynamics of consistently imaged pre-hydrolysis cell walls over time obtained using AIMTrack’s propagation strategy, including drift correction. (D) Normalized enzyme-driven AFI dynamics obtained using the AIMTrack pipeline, corrected for photobleaching.

To further characterize the autofluorescence intensity evolution during hydrolysis, temporal evolution of skewness and kurtosis of the autofluorescence intensity distributions were quantified as as measures of asymmetry, tailedness, and peakedness (Fig. S5). In the control dataset, both skewness and kurtosis remained consistently low and stable after an initial decrease. The hydrolysis dataset exhibited a small initial reduction in skewness, followed by a general upward trend, with kurtosis showing a similar trend. These results indicate that during enzymatic hydrolysis, the distributions shift away from nearly symmetrical distributions and progressively develop pronounced asymmetry and heavier tails. These results are inline with previous results reporting distinct autofluorescence distribution signatures during enzymatic hydrolysis [26]. No clear differences in skewness or kurtosis dynamics between datasets collected at 15 and 30 FPU/g biomass enzyme loadings. However, both skewness and kurtosis dynamics in the hydrolysis dataset were clearly and markedly different from the control dataset.

The autofluorescence intensity decrease observed in control datasets is due to photobleaching, a phenomenon that reduces the fluorescent signal independent of enzyme-driven biochemical changes. To accurately quantify AFI dynamics resulting exclusively from enzymatic deconstruction, the AFI dynamics observed in control datasets were used to correct those from hydrolysis datasets, by assuming that the losses attributable to enzymatic deconstruction and photobleaching combine multiplicatively [54]. The correction involved scaling the AFI decline in hydrolysis datasets by the relative AFI reduction observed in the control dataset. Therefore, at each time point, the autofluorescence intensity computed in the hydrolysis sample was divided by the corresponding reduction factor measured in the control dataset. This correction enabled isolation of the AFI loss exclusively due to enzymatic action, hereafter referred to as the enzyme-driven AFI dynamics, which exhibited a consistent, monotonic decrease over time and a dependence on enzyme loading, with higher loadings producing more pronounced reductions.

### 2.4 Enzyme-driven autofluorescence intensity dynamics are strongly correlated with conversion yields during enzymatic hydrolysis

Following photobleaching correction, AFI dynamics attributable solely to enzymatic hydrolysis exhibited a sustained, enzyme-loading-dependent monotonic decline over the 24 h period (Fig. 3.D). This clear monotonicity and dose dependence suggested that AFI dynamics could be biochemically linked to hydrolysis efficiency, motivating a direct comparison with cell wall conversion yields. Therefore, to assess the biochemical relevance of the enzyme-induced AFI dynamics, and to investigate the relationship between AFI changes and the cell wall conversion dynamics, Pearson correlation analysis was performed over 24 h period between enzyme-driven AFI dynamics and conversion yields of cell wall-derived sugars. The correlation analysis was performed separately for individual and grouped cell-wall derived sugars to determine the strength and direction of the relationship (Fig. 4).

**Fig. 4:**
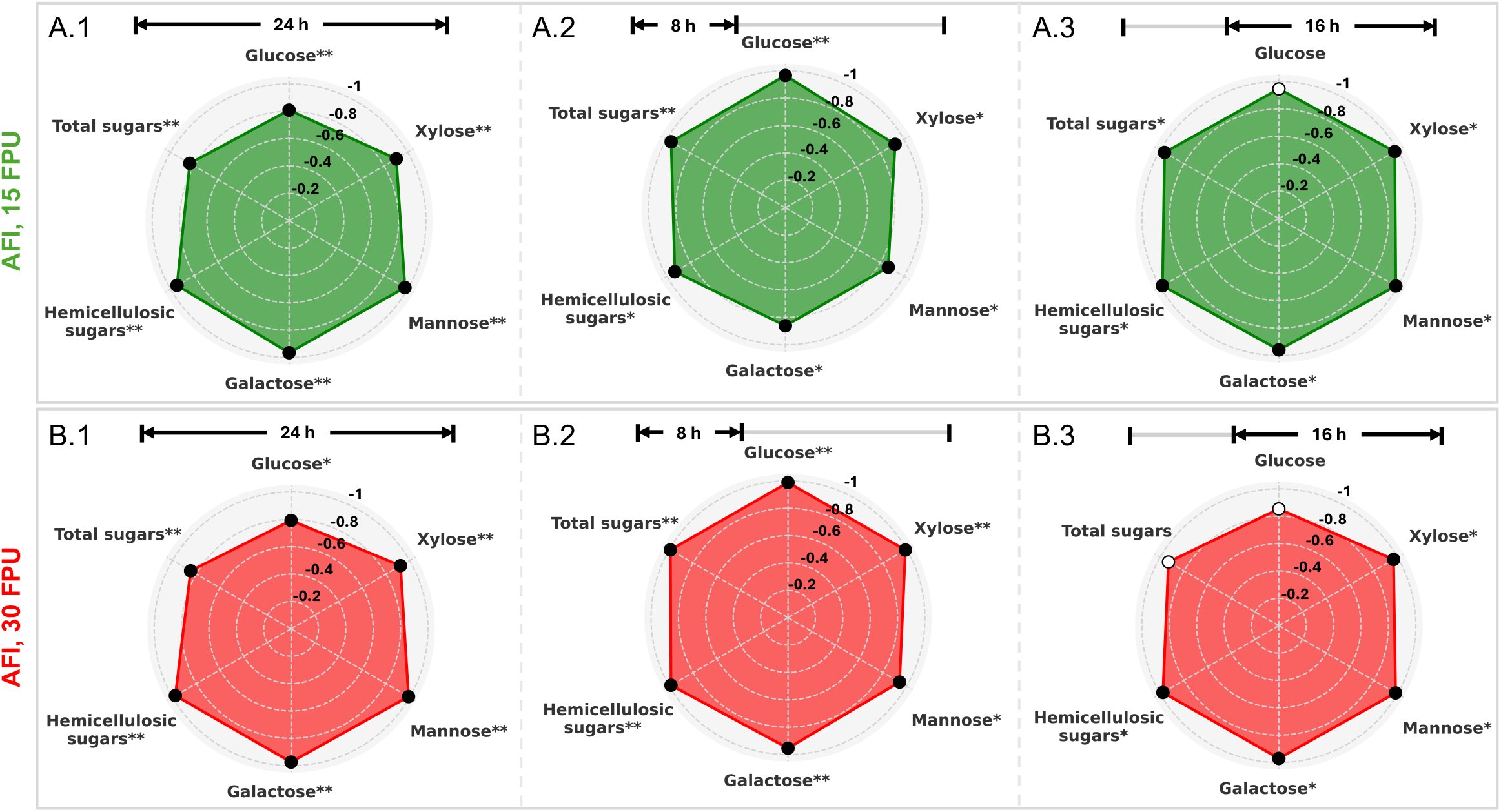
Correlation analysis between enzyme-driven autofluorescence intensity (AFI) dynamics and sugar conversion yield dynamics. (A.1–3) Pearson correlation coefficients between enzyme-driven AFI dynamics and conversion yield dynamics of glucose, xylose, mannose, galactose, combined hemicellulosoic sugars, and total sugar conversion yield dynamics at an enzyme loading of 15 FPU/g biomass, computed over the entire 24 h period (A.1), the initial 8 h (A.2), and the subsequent 16 h (A.3). (B.1–3) Corresponding Pearson correlation coefficients at an enzyme loading of 30 FPU/g biomass, evaluated over 24 h (B.1), the first 8 h (B.2), and the final 16 h (B.3). Statistically significant correlations are denoted by solid black bullets and asterisks: (* *p*-value *<* 0.05, ** *p*-value *<* 0.005). Non-significant correlations are indicated by unfilled bullets and absence of asterisks. Left-margin labels indicate the parameter correlated with conversion yields together with enzyme loading.

At 15 FPU/g biomass enzyme loading, glucose conversion yields showed strong negative correlation with enzyme-driven AFI dynamics (*r* = −0.81, *p*-value *<* 0.005). The predominant hemicellulose-derived sugars, xylose, mannose, and galactose, showed even stronger negative correlations, with *r* = −0.9, *r* = −0.98, and *r* = −0.96 respectively (all *p*-values *<* 0.005). When all hemicellulosic sugars were considered collectively, a very strong negative correlation was observed with *r* = −0.95 (*p*-value *<* 0.005). Grouping these hemicellulose-derived sugars with glucose (represented as “total sugars” in the Fig. 4) resulted again in a strong negative correlation coefficient of −0.84 (*p*-value *<* 0.005) (Fig. 4.A.1). Similarly, in the dataset collected with enzyme loading of 30 FPU/g biomass, the correlation between enzyme-driven AFI dynamics and glucose conversion yields remained strong with *r* = −0.79 (p-value *<* 0.05). Xylose, mannose, and galactose again showed very strong negative correlations with *r* = −0.92, *r* = −0.99, and *r* = −0.97 respectively (all *p*-value *<* 0.005). Grouped hemicellulose-derived sugars, exhibited a nearly perfect correlation with *r* = −0.98 (*p*-value *<* 0.005). Total sugars (hemicellulosic sugars combined with glucose), showed a lower absolute correlation coefficient of 0.85, which remained nonetheless statistically significant and strong (*p*-value *<* 0.005) (Fig. 4.B.1). A Spearman correlation analysis was also performed which revealed near-perfect to perfect monotonic associations between enzyme-driven AFI dynamics and conversion yields across all enzyme loadings and sugar types. Taken togteher, these strong inverse relationships between the enzyme-driven AFI dynamics and dynamics of conversion yields indicated that autofluorescence intensity reduction closely tracks enzymatic hydrolysis of cell wall polysaccharides into fermentable simple sugars and AFI dynamics can serve as a robust real-time proxy for assessing enzymatic hydrolysis efficiency, and most notably, for capturing the dynamics of cell wall–derived sugar conversion yields.

### 2.5 Cell wall enzymatic hydrolysis exhibits time- and sugar-type-specific correlation patterns with enzyme-driven autofluorescence intensity dynamics

The correlation analysis over the whole 24 h period provided a broad overview of the relationship between conversion yields and enzyme-driven AFI. However, this can mask phase-specific variations in the dynamics. Notably, some sugar conversion yields, particularly glucose, exhibited distinct stages, with an initial phase of rapid production during the early hours, followed by a plateau (Fig. S2). To investigate phase-specific relationships between enzyme-driven AFI changes and the conversion yields dynamics, the 24 h reaction time was divided into two intervals: the initial 8 h and the last 16 h. Separate correlation analyses were conducted for these intervals to investigate the temporal variations in the relationships between enzyme-driven AFI dynamics and sugar yield dynamics (Fig. 4.A.2, A.3, B.2, B.3).

During the initial 8 h at 15 FPU/g biomass, glucose conversion yields showed very strong negative correlation with enzyme-driven AFI dynamics with *r* = −0.97 (*p*-value *<* 0.005) (Fig. 4.A.2). Xylose, mannose, galactose, and grouped hemicellulosic sugars exhibited a strong negative correlation with *r* = −0.93, −0.87, −0.86, and −0.93 respectively with *p*-value *<* 0.05. Grouping hemicellulose-derived sugars with glucose resulted in a very strong negative correlation with *r* = −0.96 and *p*-value *<*0.005. At enzyme loading of 30 FPU/g biomass, during the first 8 h, the correlations were even stronger and, in most cases, approached near-perfect levels (Fig. 4.B.2). Glucose conversion yields exhibited nearly perfect negative correlation with enzyme-driven AFI dynamics with *r* = −0.99 (*p*-value *<* 0.005). Xylose, mannose, galactose and grouped hemicellulosic sugars, also demonstrated very strong to near-perfect negative correlations with *r* = −0.99, −0.94, −0.95, and −0.99 respectively (all *p*-value *<* 0.005, except mannose with *p*-value *<* 0.05). When glucose was grouped with hemicellulosic sugars, the correlation remained exceptionally strong at −0.99 (*p*-value *<* 0.005).

Over the last 16 h at 15 and 30 FPU/g biomass, glucose did not exhibit statistically significant correlations (*p*-value *>* 0.05) (Fig. 4.A.3, B.3). In contrast, xylose, mannose, and galactose showed very strong correlations with *r* = −0.98, −0.99, and −0.96 at 15 FPU/g biomass and *r* = −0.97, −0.98, and −0.98 at 30 FPU/g biomass, respectively (all *p*-values *<* 0.05). Grouped hemicellulosoic sugars, showed nearly perfect correlations with *r* = 0.98 and *p*-value *<* 0.05 for both enzyme loadings. The total sugars showed very strong correlations with *r* = 0.96 (*p*-value *<* 0.05) at 15 FPU/g biomass, without a significant correlation (*p*-value *>* 0.05) at 30 FPU/g biomass.

Overall, phase-specific analysis revealed that during the initial 8 h, glucose showed nearly perfect correlations, whereas in the final 16 h, glucose correlations became statistically insignificant, while hemicellulosic sugars maintained very strong correlations.

### 2.6 Temporal and sugar-specific correlations link cell wall morphological descriptors to enzymatic hydrolysis yields dynamics

Following the analysis of the relationship between enzyme-driven AFI dynamics and sugar conversion yields, the association between cell wall morphodynamics and conversion efficiency dynamics was examined. Two morphological descriptors were selected for this analysis: cell wall volume and cell-scale accessible surface area (ASA) (Fig. 5). Unlike cellulose-specific accessibility measurements reported in previous studies, the accessible surface area in this work represents the total cell-scale surface area of individual cell walls, computed from segmentations of time-lapse images generated by AIMTrack. Hereafter, ASA refers specifically to cell-scale accessible surface area.

**Fig. 5:**
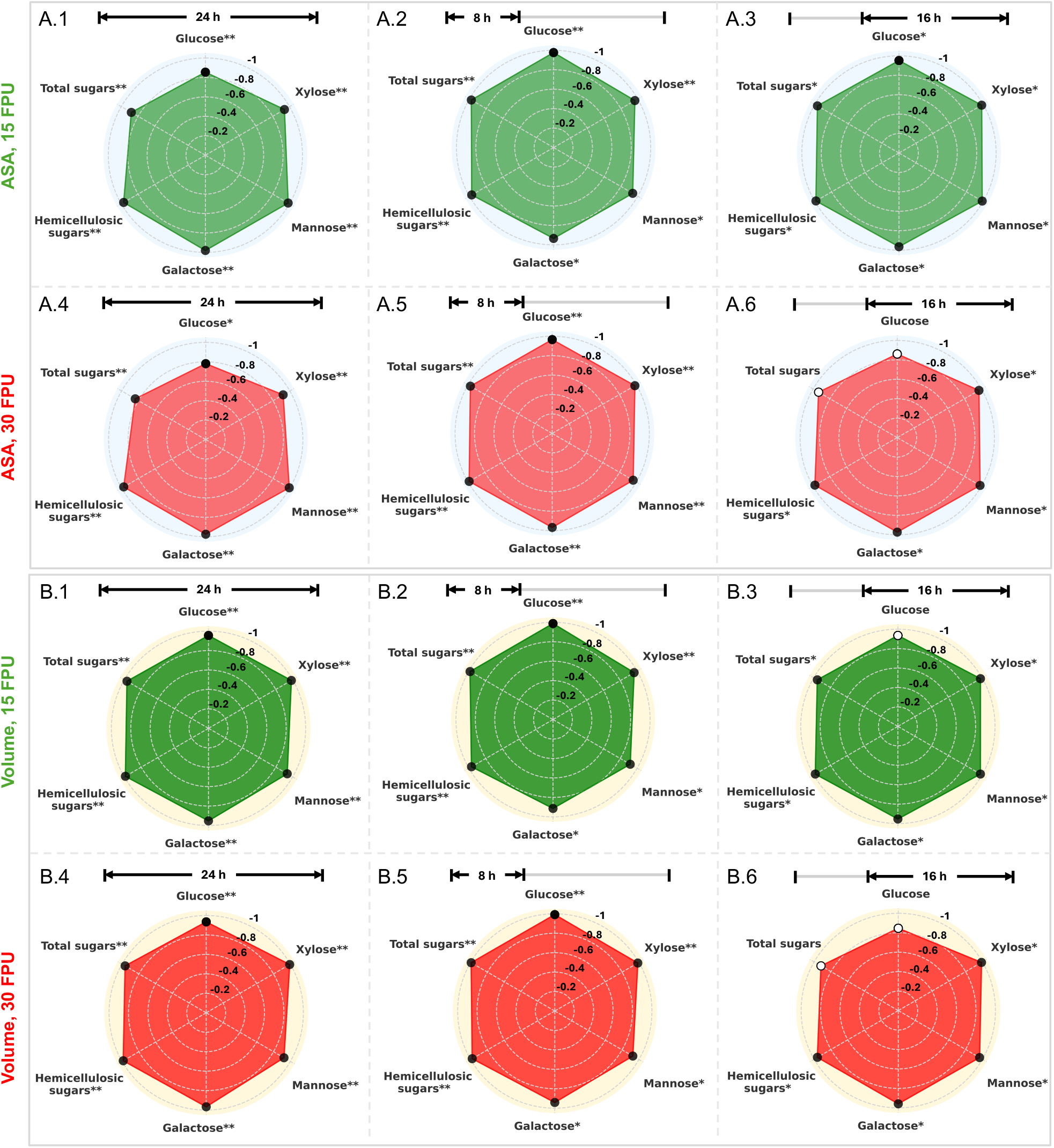
Correlation between cell wall volume and accessible surface area (ASA) dynamics and monosaccharide conversion yields. (A.1–6) Pearson correlation coefficients between normalized average ASA dynamics and sugar conversion yield dynamics at enzyme loadings of 15 FPU/g biomass (A.1–3) and 30 FPU/g biomass (A.4–6). Correlations are computed over the entire 24 h period (A.1, A.4), the initial 8 h (A.2, A.5), and the subsequent 16 h (A.3, A.6). (B.1–6) Corresponding Pearson correlation coefficients between normalized average volume dynamics and sugar conversion yields at enzyme loadings of 15 FPU/g biomass (B.1–3) and 30 FPU/g biomass (B.4–6). Correlations are computed over the entire 24 h period (B.1, B.4), the initial 8 h (B.2, B.5), and the subsequent 16 h (B.3, B.6). Statistically significant correlations are denoted by solid black bullets and asterisks: (* *p*-value *<* 0.05, ** *p*-value *<* 0.005). Non-significant correlations are indicated by unfilled bullets and absence of asterisks. Left-margin labels indicate the parameters correlated with conversion yields together with enzyme loading.

This analysis revealed strong and significant negative correlations between dynamics of conversion yields and temporal evolution of both normalized average cell wall volume and ASA, over a period of 24 h (Fig. 5.A.1, A.4 (ASA), B.1, B.4 (volume)), with all *p*-values *<* 0.005 except between ASA and glucose at 30 FPU/g biomass, where *p*-value *<* 0.05. At 15 FPU/g biomass, ASA was correlated with individual and grouped sugars (coefficients ranging from −0.85 to −0.98), and cell wall volume showed even stronger correlations (coefficients ranging from −0.94 to −0.99). At 30 FPU/g biomass, ASA correlations ranged from −0.78 to −0.99 and volume correlation ranged from −0.92 to −0.99.

To assess how relationship may evolve over time between both cell wall volume and cell-scale ASA and dynamics of conversion yields, the 24 h hydrolysis period was divided into two phases: an early phase (0–8 h) and a late phase (8–24 h), following the phase-specific analysis previously applied to AFI dynamics and conversion yield dynamics. Pearson correlation analyses were performed separately for early and late phases. The results showed that during the initial 8 h, strong negative correlations were observed between dynamics of conversion yields and the dynamics of normalized average cell wall volume and ASA at both enzyme loadings. At 15 FPU/g biomass, correlation coefficients ranged from –0.91 to –0.99 for volume (Fig. 5.B.2) and –0.93 to –0.98 for ASA (Fig. 5.A.2). Similarly, at 30 FPU/g biomass, correlations ranged from –0.93 to –0.99 for volume and ranged from –0.96 to –0.99 (Fig. 5.B.5) for ASA (Fig. 5.A.5). These results indicate a consistent and strong inverse relationship between early-phase structural degradation and conversion yields for all sugar types. All correlations with glucose, xylose, hemicellulosic sugars, and total sugars were statistically significant with *p*-values *<* 0.005. For mannose and galactose, *p*-values were consistently *<* 0.05, except for ASA at 30 FPU, where the *p*-values were also *<* 0.005. During the subsequent 16-h phase, negative correlations between sugar yields and cell wall structural dynamics remained generally strong but exhibited notable differences in statistical significance based on sugar type and enzyme loading. Notably, correlations involving glucose generally lost statistical significance (*p*-value *>* 0.05). A similar loss of significance was observed at the higher enzyme loading (30 FPU/g biomass) between total sugars and both normalized average volume and ASA dynamics. In contrast, correlations with individual and grouped hemicellulosic sugars remained consistently strong and statistically significant across both enzyme loadings and morphological descriptors (*p*-value*<* 0.05).

To further characterize enzyme-induced cell wall structural changes, dynamics of average cell wall deformation between the pre-hydrolysis image and each subsequent time point were quantified (Fig. 6). The results revealed minor deformations in the control datasets. In contrast, hydrolysis dataset showed a time-dependent, monotonic increase in deformation that scaled with enzyme loading: the hydrolysis dataset at 30 FPU/g biomass exhibited the greatest deformation with steep increase after 8 h, remarkably exceeding that observed at hydrolysis dataset at 15 FPU/g biomass. A further correlation analysis revealed cell wall deformation being tightly linked to normalized average volume and ASA dynamics during enzymatic hydrolysis. At 15 FPU/g biomass, the correlation coefficients were similarly high for ASA (*r* = –0.96, *p*-value *<* 0.005) and volume (*r* = –0.97, *p*-value *<* 0.005), indicating nearly equivalent relationships. In contrast, at 30 FPU/g biomass, ASA showed a stronger negative correlation with deformation (*r* = –0.95, *p*-value *<* 0.005) compared to volume (*r* = –0.82, *p*-value *<* 0.005), suggesting that at higher enzyme loadings, changes in ASA are more tightly associated with structural deformation than changes in volume.

**Fig. 6:**
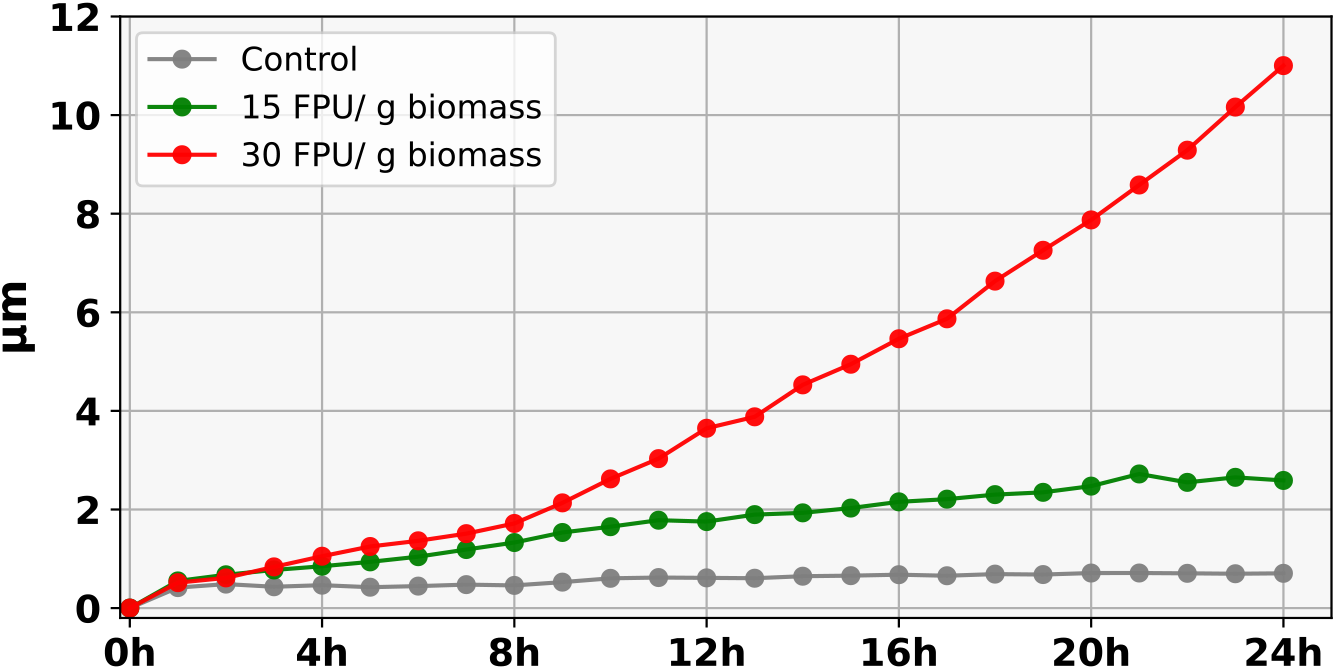
Cell wall deformation dynamics during enzymatic hydrolysis for control and hydrolysis datasets.

## 3 Discussion

A key contribution of this study is the development of the AIMTrack pipeline, which enables automated analysis of time-lapse 3D acquisitions of plant cell wall enzymatic deconstruction. AIMTrack provides accurate quantification of autofluorescence intensity dynamics together with dynamics of morphological descriptors of the cell wall, including volume, cell-scale ASA, and deformation. AIMTrack, includes a novel efficient adaptive compensation strategy for sample drift and deformation that adapts to the extent of deconstruction. Sample drift is a common challenge in imaging that should be appropriately addressed to ensure reliability and accuracy of downstream quantitative analyses. Most drift correction methods have been developed for and applied to nanoscale imaging such as single-molecule localization microscopy [4, 13, 34]. In microscale imaging, drift compensation remains challenging, particularly in studies of plant cell wall enzymatic deconstruction where different approaches have been proposed [26, 49]. AIMTrack addresses the limitations of the existing drift compensation strategies such as global, local, or fixed-size clustering, by introducing an innovative adaptive clustering approach that dynamically adjusts cluster sizes based on the extent of deconstruction, employing larger clusters during slow hydrolysis periods and smaller ones during periods of significant structural changes. This adaptive clustering strategy is an enhanced divide-and-conquer strategy with a context-aware partitioning mechanism, where the cluster boundaries are modulated by progression of enzymatic hydrolysis. Given the intrinsic temporal heterogeneity of enzymatic hydrolysis dynamics, this data-driven adaptivity is essential to achieve accurate characterization. Subsequently, AIMTrack provides robust and precise quantification of autofluorescence intensity and cell wall morphological descriptors extracted from time-lapse images. AIMTrack is also highly versatile, applicable to time-lapse images across diverse biomass species, pretreatment types, and enzymatic cocktails. These advantages position AIMTrack as a broadly applicable pipeline for microscale structural analyses of lignocellulosic biomass during enzymatic hydrolysis.

Application of AIMTrack to time-lapse images of pretreated spruce wood enzymatic hydrolysis, revealed strong and consistent negative correlations between enzyme-driven AFI dynamics and dynamics of conversion yields of individual and grouped sugars, namely glucose and hemicellulose-derived sugars, particularly xylose, mannose, and galactose over 24 h. This strong inverse relationship indicated that enzyme-driven AFI dynamics closely tracks enzymatic hydrolysis yield dynamics of cell wall polysaccharides into fermentable simple sugars. Wood cell wall autofluorescence primarily originates from phenolic compounds that constitute lignin [35], and its intensity depends on a range of chemical and physical factors, including fluorophore content, inter-linkage types, and interactions with other cell wall polymers [15]. These dependencies suggest that enzymatic deconstruction not only alters polysaccharide content but also leads to substantial modifications in the lignin environment, as indirectly reflected by the progressive decrease in autofluorescence intensity and directly supported by the release of sugars from hydrolyzed polysaccharides. While previous work showed that initial cell wall autofluorescence intensity correlates with final released glucose concentration after prolonged hydrolysis [3], this study takes a different dynamic approach. By investigating the temporal dynamics of enzymatic deconstruction, real-time correlations of conversion yields during hydrolysis with both cell wall morphodynamics and autofluorescence intensity dynamics are revealed.

More importantly, phase-specific analysis revealed distinct temporal and sugar-type-dependent patterns in the correlation between enzyme-induced AFI dynamics and sugar conversion yields. During the initial 8 h, dynamics of glucose conversion yields, which mostly originated from cellulose (representing nearly 60% of the sugars (Fig. S1)) exhibited near-perfect negative correlations with fluorescence loss, indicating rapid early-stage hydrolysis. In contrast, over the final 16 h, glucose correlations became statistically insignificant while hemicellulose-derived sugars—particularly xylose, mannose, and galactose—maintained strong correlations, independently of enzyme loading. These results reveal temporally resolved deconstruction phases and substrate-specific enzymatic activity dynamics. Given that chlorite-based pretreatment leads to a significant removal of lignin (nearly 63 %), cellulose fibers become much more accessible, as previously illustrated for instance for poplar wood [14]. Discrepancy between the first 8-h phase and the following phase highlights the phenomenon known as cellulosic hydrolysis slowdown [2], mostly attributed to the irreversible deposition of non-cellulosic species (either reaction side products or denatured enzymes, or both) on the cellulosic substrate surface [29]. Conversely, hemicellulosic sugars conversion appears consistent over hydrolysis time, probably because enzymatic activities are in limited amounts given the relative hemicellulose content in comparison to cellulose. These results suggest that autofluorescence intensity dynamics allows tracking distinct phases of cell wall enzymatic deconstruction and offers a kinetic window into cell wall deconstruction, which should be taken into account when used for predictive computational modeling of lignocellulosic biomass enzymatic deconstruction and conversion optimization strategies such as enzyme cocktail composition and tailored enzyme application strategies.

Finally, our findings also revealed enzyme mediated cell wall structural changes including cell wall volume, ASA, and deformation reflecting enzymatic deconstruction efficiency dynamics. Strong, statistically significant negative correlations were observed across enzyme loadings (15 and 30 FPU/g biomass) between dynamics of individual and grouped sugar conversion yield and dynamics of cell wall volume and ASA, over 24 h. Temporal analysis revealed that this inverse relationship was most pronounced during the early hydrolysis phase (0–8 h), where correlations remained consistently strong and significant across all sugars and enzyme loadings. In contrast, during the late phase (8–24 h), correlations involving glucose and total sugars weakened or lost significance, while those for hemicellulosic sugars remained robust. Quantification of cell wall deformation further supported these findings, showing monotonic increases with enzyme loading and time. Deformation dynamics were strongly correlated with both ASA and volume, with ASA showing a stronger association with deformation than volume. Overall, these results indicate that enzymatic hydrolysis progressively compromises cell wall structural integrity, leading to reductions in both volume and cell-scale ASA, and resulting in increased mechanical deformability. In return, the observed correlations with sugar conversion dynamics underscore the predictive potential of quantitative morphology-based analyses for understanding and predicting hydrolytic efficiency and conversion dynamics.

Previous studies and models of enzymatic cellulose digestion based on quantitative AFM data, revealed formation of elongated cracks on the substrate surface upon cellulase treatment, highlighting localized structural changes during hydrolysis [18]. However, these findings are limited in scope due to their reliance on purified substrates lacking native cell wall complexity. Similarly, particle-scale imaging of hardwood and switchgrass hydrolysis revealed progressive shape alterations over time [33], aligning conceptually with our observations. Yet, these analyses were conducted at the 10–100 µm scale using pure cellulase cocktails, precluding identification of specific structural markers at the cell wall level. Some efforts have been done for providing 3D analysis of cell wall after pretreatment of spruce [42] or during enzymatic hydrolysis of wheat straw by X-ray tomography, but without a detailed investigation of the relationship between structural changes, biochemical composition, and deconstruction dynamics. [8].

In contrast, and to the best of our knowledge, this study is the first to establish a direct and quantitative relationship between dynamics of conversion yields, cell wall autofluorescence intensity, and morphological descriptors at the cell wall scale. Furthermore, it characterizes the temporal dynamics of these interrelated parameters under conditions of extensive substrate conversion, an essential context for optimizing hydrolytic efficiency. The results align with prior work demonstrating the relevance of microscale structural parameters to investigate the enzymatic deconstruction efficiency under limited conversion conditions in poplar wood [49].

Overall, this study establishes a robust, quantitative, and nondestructive approach for assessing plant cell wall conversion efficiency, eliminating the need for extensive sampling and labor-intensive chemical assays. By validating cell wall morphological descriptors and autofluorescence intensity as accurate predictive real-time biomarkers of conversion yields, the findings provide a powerful tool for evaluation of enzymatic deconstruction efficiency and dynamics of cell wall sugars yields. Our approach offers substantial potential to advance the development and optimization of biotechnological strategies for efficient plant biomass deconstruction and conversion.

## 4 Methods

### 4.1 Pretreatment and composition analysis

Spruce (*Picea abies*) samples, supplied by INRAE, Champenoux, Grand-Est, France, were first cut into fragments measuring 0.5 × 0.5 × 1 cm. Sample fragments were then pretreated using sodium chlorite to achieve partial delignification. Specifically, the samples were immersed in eight consecutive treatment baths, each lasting 1 h at 70 °C, in a solution composed of 1.25 g of sodium chlorite, acetic acid, and 40 mL of water [22].

To determine lignin content and polysaccharides composition, the 3 mm end segments of raw and pretreated sample fragments, selected for consistency with the subsequent microscopy acquisitions, were first ground. Lignin content was measured by acetyl bromide assay in triplicate [41]. Sugar composition was determined in triplicate following acid hydrolysis. Hydrolysates were analysed by high-pressure anion exchange chromatography (HPAEC - Dionex ICS5000, ThermoFisher, USA), equipped with Carbopac PA1 column (4×25mm) and precolumn, using gradient elution with sodium hydroxyde and sodium hydrogenocarbonate and detection by pulsed amperometry [7].

### 4.2 4D (3D + time) confocal fluorescence microscopy imaging

To reduce sample spatial heterogeneity that could be introduced by pretreatment differential impact, only the terminal 3 mm regions along the 1 cm longitudinal axis of the pretreated samples were used for sectioning. These extremities were selected as they were consistently and thoroughly exposed during the pretreatment. Sections of 40 µm thickness were then prepared from these specific regions using a microtome (HM340E, Thermo Scientific, Waltham, USA) equipped with a disposable blade. A sample section was then placed onto a glass slide and a Gene Frame was positioned around it. 60 µL of acetate buffer (50 mM, pH 5) was then added within the Gene Frame and incubated for 30 minutes. Another sample section was weighed with a precision balance. A cellulolytic enzyme cocktail (IFPEN, Rueil-Malmaison, France), was diluted with acetate buffer, to reach 15, and 30 FPU/g biomass. At the end of the 30-minute incubation, filter paper was used to remove the buffer within the Gene Frame. A volume of 60 µL of enzyme solution (or acetate buffer in case of control dataset) was then added into the frame, which was subsequently sealed with a coverslip. The prepared slide was placed onto the stage of a confocal microscope (Olympus Fluoview FV3000, Tokyo, Japan) within a temperature-controlled incubation chamber (OKOlab, Italy) maintained at 50°C [61]. Time-lapse imaging was then performed by acquiring four z-stacks at 1 h intervals for a period of 24 h. Imaging was performed using a 405nm excitation laser set to 10% power, with fluorescence emission detection in 415-515nm range. Scanning was conducted at 4µs/pixel with a resolution of 512×512 pixels, and the gain was set to 600V. The confocal pinhole aperture was adjusted to 50µm. For time-lapse imaging, a 20× objective lens was used with a 2× digital zoom. Z-stacks, each containing 400 sequential optical sections, were acquired with a step size of 0.3 µm along the Z-axis.

### 4.3 Conversion yields quantification during enzymatic hydrolysis

Sample sections, prepared as described previously in section 4.1, were placed into individual 200 µL mini-reactors, to which 60 µL of enzyme solution (or acetate buffer to collect control dataset) was added. Each mini-reactor tube was closed with a lid and wrapped with some Parafilm to ensure effective prevention of evaporation. Experiments were performed in triplicate mini-recator tubes for each enzyme loading. Each tube was then placed in the incubation chamber stabilized at 50°C. Sampling of the reaction mix into a eppendorf tube was performed at 0h, 15 min, 1 h, 2 h, 4 h, 8 h, 12 h, 16 h, 20 h, 24 h after incubation launch. Hydrolysates were then boiled for 5 minutes, then briefly centrifuged and stored at −20°C prior to analysis. For quantification of released sugars, 30 µL aliquots of enzymatic hydrolysates were mixed with an internal standard and diluted prior to analysis by High-Performance Anion Exchange Chromatography (HPAEC) using a Dionex ICS-5000 system (Thermo Fisher Scientific, USA).

### 4.4 4D image processing

Let 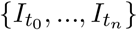 denote the set of time-lapse 3D images of plant cell wall deconstruction, where 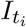 represents the *i*-th 3D confocal image (z-stack) acquired at time *t*_*i*_, with 0 ≤ *i* ≤ *n*, and 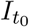 and 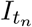 denoting the pre-hydrolysis and final acquisitions respectively. The time-lapse 3D images represent cell wall autofluorescence intensity images. AIMTrack is devised to analyse time-lapse 3D confocal images through a workflow involving the following steps:

- Adaptive temporal clustering of time-lapse images
- Drift and deformation compensation
- Tracking cell walls in time-lapse autofluorescence intensity images using a propagation strategy
- Identification of individual cell walls

In the following, a detailed explanation of each step is provided.

#### 4.4.1 Adaptive temporal clustering of 3D time-lapse images

Direct registration of temporally close images with conventional registration algorithms can be performed with reasonable accuracy. However, direct registration often fails when applied to temporally distant images, in particular under extensive deconstruction conditions, leading to excessive deformation of the floating image [26]. Indirect registration approaches [26], address this issue by dividing time-lapse images into clusters of fixed size, constraining the registration time-frame to the images within clusters and composing the resulting transformations to register temporally distant images. This fixed size clustering has the following limitations: (1) the cluster size needs to be determined manually which is laborious and time-consuming, as it requires repeated trials and visual inspection by an expert to determine the cluster size. (2) fixed-size clusters implicitly assume a uniform rate of deconstruction throughout the hydrolysis process. Consequently, the use of a fixed cluster size requires setting the cluster size according to the fastest deconstruction stage, leading to unnecessary subdivision of image sequences during stages with limited deconstruction. This over-clustering compromises the quality of results and downstream quantifications [26]. AIMTrack addresses these limitations by defining clusters as maximal contiguous sequences of time points within which the cell wall deformation remains below a specified threshold. In doing so, AIMTrack automatizes the clustering process and reduces the number of clusters required to achieve accurate registration compared to fixed-size clustering, which can excessively fragments timelapse sequences, and introduce accumulation of registration errors generated by an increased frequency of transformation compositions.

To achieve this, AIMTrack divides the time-lapse images into *v* sequential clusters of variable sizes, denoted by *C*_*q*_, *q* ∈ {0, 1, …, *v* − 1}. Each cluster shares a single image with its preceding cluster such that its first image is the last image of the previous cluster (Fig. 1). Each cluster *C*_*q*_ consists of *m*_*q*_ + 1 successive images. The clusters are defined as follows:

- The first cluster, *C*_0_, consists of the first *m*_0_ + 1 images:

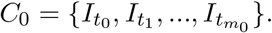
- For *q* ≥ 1, the cluster *q* is defined as :

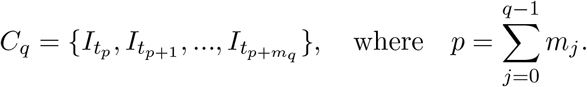

Cluster sizes, *m*_*q*_, are determined adaptively according to the extent of deconstruction over specific intervals, defining cluster boundaries based on deformation that reflect the extent of structural change over time. To accomplish this, AIMTrack initiates clustering from the pre-hydrolysis image and assesses deformation between the first image of the cluster (here pre-hydrolysis image) and subsequent hydrolysis-phase images using registration both in forward and backward directions. This deformation is quantified as a vector field resulting from nonlinear registration, performed after sequentially applying rigid and affine registrations [48]. The nonlinear registration computes the residual deformation as a vector displacement field. The block matching framework is used for the registrations [40, 44]. Deformation magnitude, defined as the largest of the means of Euclidean norms of vectors comprising these vector fields derived from forward and backward registrations, is then computed. Cluster expansion proceeds progressively, integrating subsequent images while the deformation magnitude does not exceed a predefined threshold value (denoted by *ϵ* and set at 4.1 for hydrolysis and control datasets, chosen by an expert as the maximum tolerable deformation). Upon exceeding this threshold, AIMTrack terminates the current cluster and initiates a new cluster starting from the last image of the preceding cluster. This iterative clustering process continues systematically until reaching the last image of the time-lapse series. The adaptive clustering ensures that intra-cluster deformations remain below the predefined threshold. Consider a given cluster, 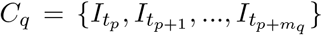 with 0 ≤ *q < v* and 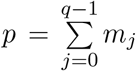. Let 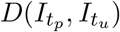 denote the deformation field magnitude. By construction, the deformation within the cluster satisfies 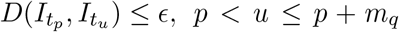, while including of the next image would exceed the threshold: 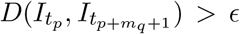 (Fig. 1).

##### Benchmarking AIMTrack adaptive clustering

To assess the accuracy and performance of the AIMTrack clustering, the clusters generated with AIMTrack were compared with expert determined clusters obtained through expert visual inspection. Expert determination of clusters involved visual inspection of the registration quality by generating an RGB image in which the tracked region of the reference image was assigned to the green channel and the registered floating image to the red channel and subsequently inspecting the overlay of the green channel and red channel with ImageJ. In the RGB image, the untracked region of the reference image due to sample drift was saved in blue channel. Cluster boundaries were manually defined at time points where misalignments appeared as color fringes which indicated registration failure. In fact, successful registrations are characterized by overlapping red and green signals which combine to produce yellow in the RGB overlay while imperfect registration creates color fringes as visible edges of pure red or pure green. These expert-defined clusters served as a benchmark for evaluating the clustering results produced by AIMTrack. The cluster time points identified by AIMTrack were compared to those annotated manually, revealing strong agreement: 213 out of 225 time points (approximately 95%) matched the expert annotations (1).

To further evaluate the performance of adaptive clustering of AIMTrack, the average cluster sizes obtained using AIMTrack were compared to fixed-size clustering strategy (Fig. 2) given that in the fixed cluster size clustering the smallest cluster size detected by AIMTrack should be used uniformly across all time points of a time-lapse sequence. Larger clusters imply fewer required successive application (or compositions) of transformations across clusters and smaller clusters result in higher number of compositions, increasing cumulative registration error [26]. This relationship between cluster size and accuracy was considered in the comparative evaluation of AIMTrack and fixed-size clustering strategies.

#### 4.4.2 Drift and deformation compensation

AIMTrack uses the pre-hydrolysis image 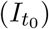 as the reference image for drift and deformation compensation. The hydrolysis-phase images are registered with pre-hydrolysis image. To achieve this, following adaptive clustering of the time-lapse images, intra-cluster bidirectional transformations are computed: in the forward direction, the first image of each cluster is registered to subsequent images within the cluster; in the backward direction, each cluster image is registered to the first image of the cluster (Fig. 1.B.2, Fig. S4.A). Each registration involves first computing a rigid transformation followed by computing an affine transformation. This is then followed by computing a non-linear transformation. The rigid, affine, and non-linear transformations are computed using the block matching framework.

To register the hydrolysis-phase image, 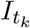, 0 *< k* ≤ *n* with the pre-hydrolysis image, the cluster *C*_*s*_ which includes 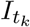 is first identified. If 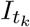 is a boundary image, 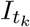 belongs to two consecutive clusters and the left one is chosen. *C*_*s*_ can be written as:

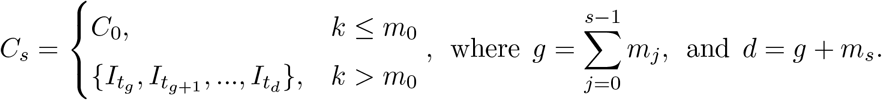

Let 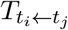 denote the transformation that registers 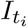 (as the floating image), with the 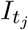 (as the reference image), for 0 ≤ *i, j* ≤ *n*. The resampled image is denoted by 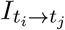 which is obtained by applying the resulting transformation to the floating image, 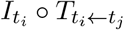.

When *k* ≤ *m*_0_, the resampled image is directly computed as

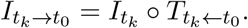

For *k > m*_0_, 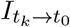 which is the resampled image, is obtained through the recursive application of transformations:

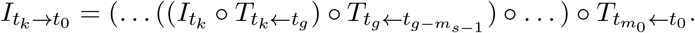

#### 4.4.3 Tracking cell walls in time-lapse autofluorescence intensity images using a propagation strategy

Following the approach described in [26], the resampled images 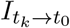 are used to detect the pairwise temporally consistent imaged regions between 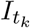 and 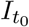, 0 *< k* ≤ *n*. For each 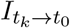, a binary image, denoted by 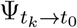, is created where voxels which map back to voxels in 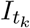 are set to 1. Therefore, 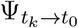 marks the subset of 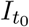 which is overlapped by 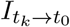. The consistently imaged region, denoted by 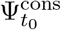, which corresponds to the subset of the sample that remains within the microscope’s field of view during time-lapse acquisitions (Fig. 1.C), is determined by computing the intersection of these masks obtained using a logical AND:

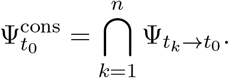

The cell walls located within this region, referred to as the pre-hydrolysis consistently imaged cell walls and denoted by 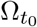, are then identified using thresholding. To track this region, temporal propagation of spatial information strategy is employed using the forward transformations to propagate 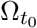. This region at time point *t*_*k*_ is denoted by 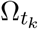, 0 *< k* ≤ *n* (Fig. 1.D, Fig. S4.B). The propagated region at time point *t*_*k*_ is denoted by 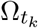, where 0 *< k* ≤ *n* and is obtained as follows:

- If *k* ≤ *m*_0_

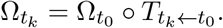
- For 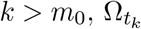 is computed using:

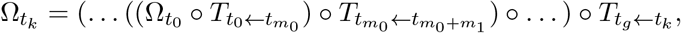

where 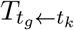 is the transformation which enables to register the left boundary image, 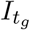, of the cluster including 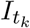, with 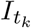. In case 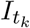 itself is a boundary image belonging to two consecutive clusters, and 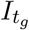 corresponds to the left boundary image of the preceding (left) cluster.

#### 4.4.4 Identification of individual cell walls

To identify and track individual cells in the time-lapse images, the pre-hydrolysis image is first segmented using the previously developed method [26] which involves computing the convex hull of the imaged sample in pre-hydrolyis image. The individual cells are then identified by spatially constraining the watershed algorithm to the region inside the convex hull, generating a segmented image denoted by 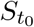. Similar to computation of 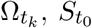 is then propagated throughout the hydrolysis (Fig. 1.E, Fig. S4.C) phase as follows:

- When *k* ≤ *m*_0_:

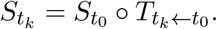
- For *k > m*_0_:

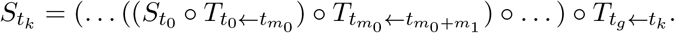

The reduced number of clusters achieved through AIMTrack’s adaptive clustering strategy, and consequently the decreased frequency of required registrations and transformation composition and application, positively influences the identification of individual cell walls in hydrolysis-phase images and quality of segmentations as previously demonstrated [26].

### 4.5 Estimation of cell-scale accessible surface area

To estimate the morphology-based accessible surface area at the cell scale (ASA), defined as the surface area of the interface between the cell walls of individual cells and the background, the Marching Cubes (MC) algorithm from the Visualization Toolkit (VTK)[55] library was employed to generate a mesh of the cell walls. This mesh was then decimated, smoothed using a Laplacian filter, and subsequently used to estimate the surface area. The decimation and smoothing steps were used to optimize the surface area estimation. Two parameters in the VTK decimation and smoothing methods were optimized to improve estimation accuracy: `TargetReduction`, which defines the level of mesh reduction (0 ≤ TargetReduction ≤ 1), and `NumberOfIterations`, which sets the maximum number of smoothing iterations. For this optimization, 100,000 three-dimensional objects in the form of cell-shaped polyhedra (right prisms with randomly generated holed polygonal bases) were generated with known surface areas. The parameters were tuned on these randomly generated objects to obtain surface area estimations with minimal mean squared error (MSE).

Due to the possibility of local minima, three criteria were used to select the optimal parameters, which are listed in the following in descending order of importance: (1) the MSE, (2) the estimation bias, and (3) the mean CPU time required to perform a single surface area estimation, including decimation and smoothing. These criteria were evaluated using a Monte Carlo approximation, which generated optimal parameter values of 0.1 for `TargetReduction` and 50 for `NumberOfIterations`.

The resolution chosen for the optimization was 80, meaning that for each generated cell-shaped polyhedron, the cell diameter corresponded to 80 times the voxel side in the horizontal (*x, y*) plane, and the cell thickness was set to half of the cell diameter. These dimensions were selected because they most closely represent the true cell dimensions observed in the pre-hydrolysis images.

To evaluate the performance of the optimal VTK parameters, 1,656 virtual cell-shaped polyhedra approximating the original true cells were generated. This generation process involved: (i) projecting the original cells onto the horizontal (*x, y*) plane, (ii) approximating the resulting two-dimensional digital objects using the `approximate_polygon` function from the *scikit-image* package, and (iii) constructing three-dimensional cell-shaped polyhedra by assigning each polygon a uniform thickness equal to the original cell wall thickness, obtained from projections onto the vertical *z* axis. Using the optimal parameters values, a mean *AD* of 0.9% and a *CV* of 1.97% were obtained for the virtual cells. The mean *AD* denotes the mean absolute deviation defined as:

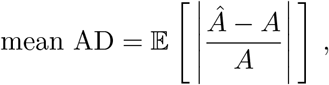

with Â denoting the estimated surface area, *A* the ground truth surface area, 𝔼[·] the expectation operator, and | · | the absolute value. *CV* denotes the coefficient of variation, defined as *σ/µ*, where *σ* is the standard deviation of the estimation errors and *µ* is their mean. These results demonstrate that using optimized parameters enable highly accurate surface area estimations for cell-shaped polyhedra closely approximating the true cells in pre-hydrolysis images. Subsequently, these parameter values were used to estimate the cell-scale ASA. The accuracy achieved in estimating surface area combined with the precision of AIMTrack’s image processing framework, underscores the overall robustness and reliability of the quantifications obtained using AIMTrack pipeline.

## Acknowledgments

We gratefully acknowledge Simon Arragain and Antoine Margeot (IFPEN, Rueil-Malmaison, France) for provision of enzymatic cocktail. We also wish to thank Juliette Floret for measuring enzymatic activity and François Gaudard for his help for preparation of the samples and measuring conversion yields. We also thank Grégoire Malandain for the block matching algorithm.

## Author contribution

S.HK., G.P., and Y.R. conceptualized and designed the research. G.P. and Y.R. supervised the project. S.HK., K.OA., M.C., A.F., G.P., and Y.R. developed the methodology. N.R., A.H., B.L., and G.P. collected time-lapse image datasets and conversion yields. S.HK., K.OA., F.SG., M.C., J.M., G.P., Y.R. analysed the data. S.HK., K.OA., G.P., and Y.R. wrote the original draft with inputs from all authors. All authors reviewed and approved the manuscript.

## Funding

This work was funded by French Research Minister aid managed by the Agence Nationale de la Recherche as part of the France 2030 investment plan though the grant reference ANR-23-PEBB-0006 (FillingGaps project) awarded to G.P. This work was also funded by Agence Nationale de la Recherche (ANR) under grant ANR-19-CE43-0010 (BIOMOD project) awarded to Y.R., and by Grand Est Region through “BIOMODEL” doctoral funding to S.HK.

## Competing interests

The authors declare no competing interests.

## Supporting Information for

The supporting information includes four figures, Fig. S1, Fig. S2, Fig. S3, and Fig. S4.

**Fig. S1:**
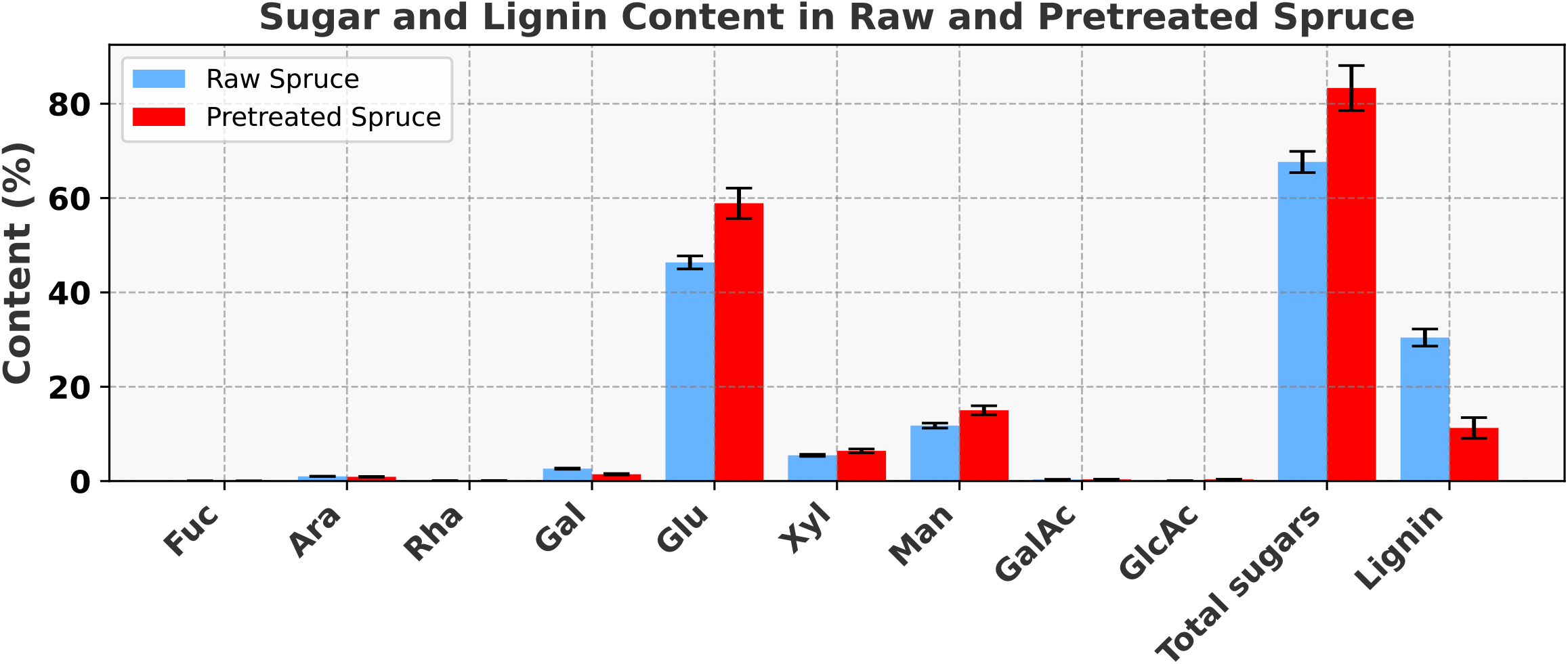
Chemical composition of raw and pretreated spruce samples.

**Fig. S2:**
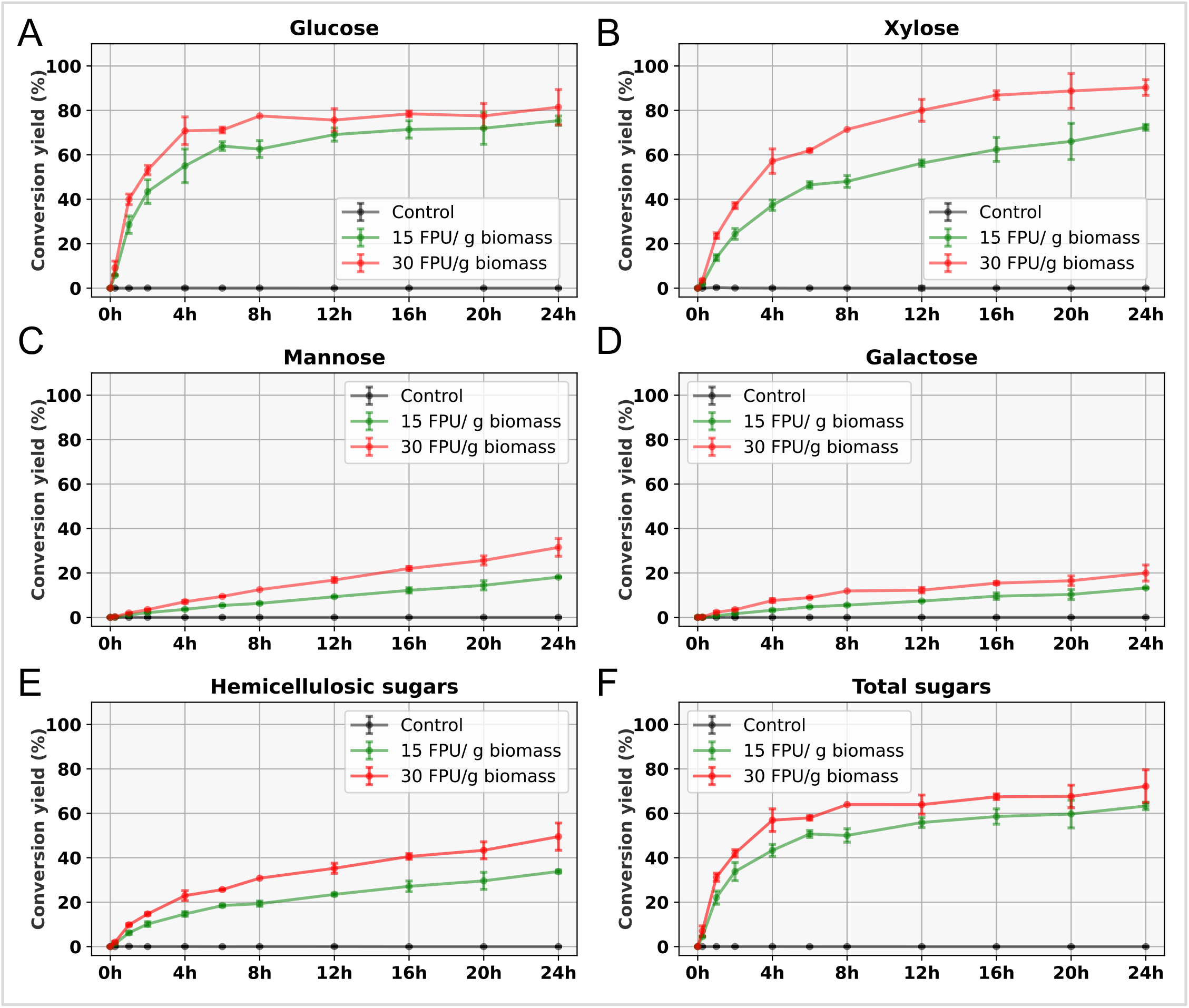
Sugar conversion yields of pretreated samples, including the control and the hydrolysis datasets with enzyme loadings of 15 and 30 FPU/g biomass. (A–D) Conversion yield dynamics for glucose, xylose, mannose, and galactose, respectively. (E) Combined hemicellulosic sugar conversion yield dynamics including mainly mannose, xylose, galactose. (H) Total sugar conversion yield dynamics which include hemicelluosic sugars and glucose.

**Fig. S3:**
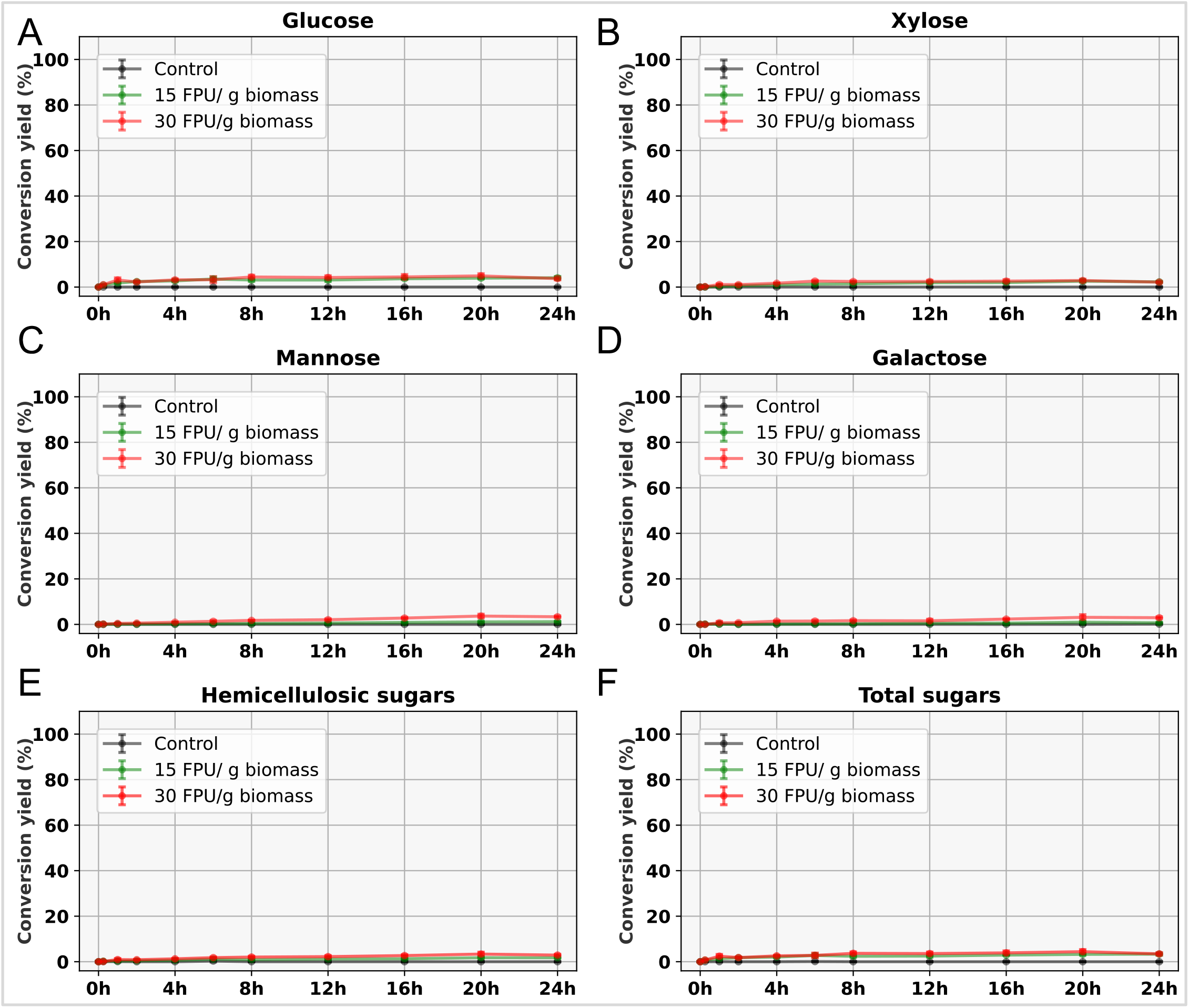
Sugar conversion yields of raw samples,, including the control and the hydrolysis datasets with enzyme loadings of 15 and 30 FPU/g biomass. (A–D) Conversion yield dynamics for glucose, xylose, mannose, and galactose, respectively. (E) Combined hemicellulosic sugar conversion yield dynamics including mainly mannose, xylose, galactose. (H) Total sugar conversion yield dynamics which include hemicelluosic sugars and glucose.

**Fig. S4:**
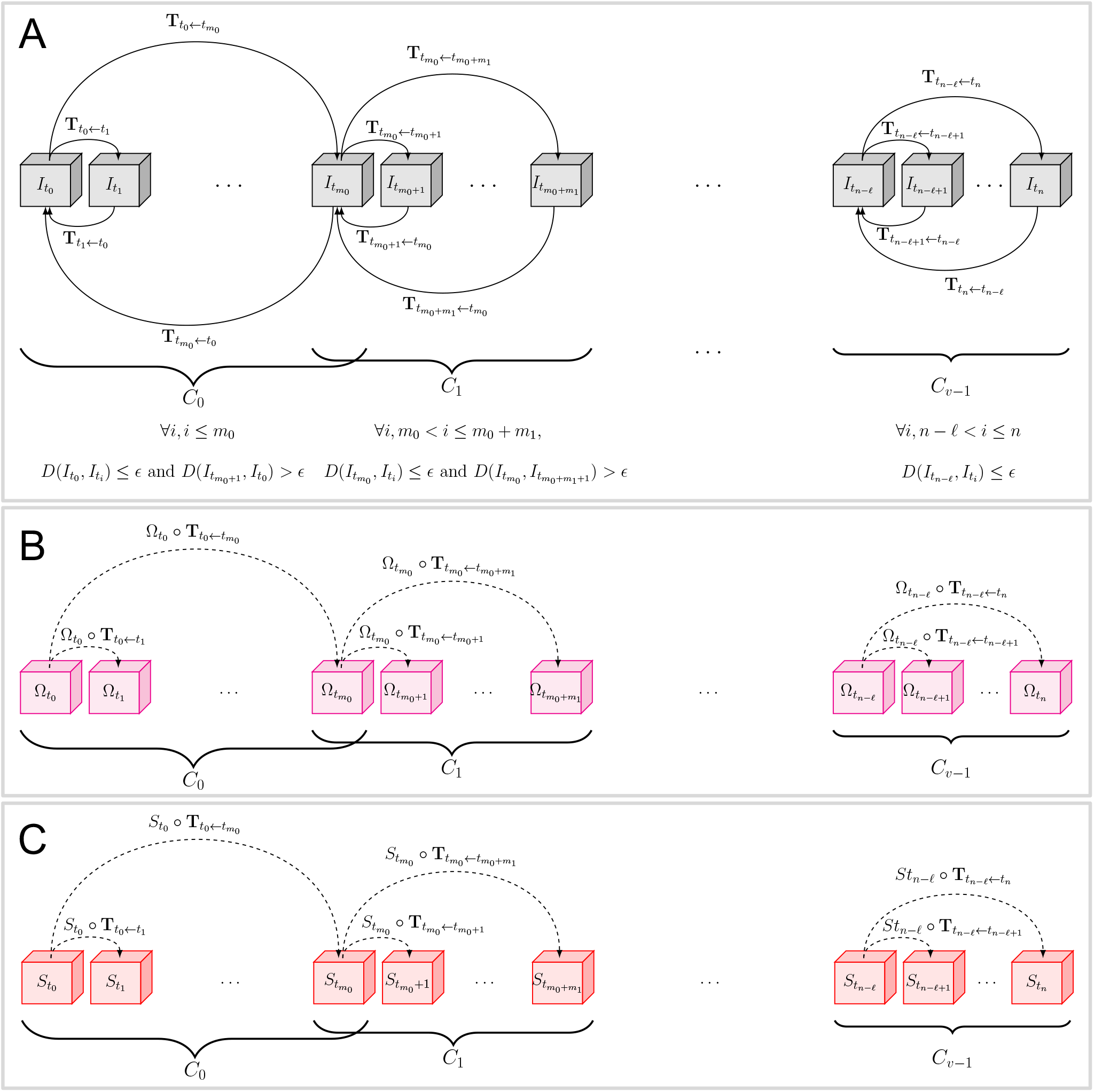
AIMTrack pipeline to track cell wall enzymatic hydrolysis. (A) Adaptive clustering of time-lapse images. 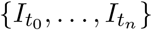 denotes the time-lapse 3D images with 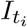 acquired at time *t*_*i*_, 0 ≤ *i* ≤ *n*. Starting from 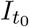, consecutive images are adaptively grouped into clusters based on deformation magnitudes. The deformation magnitude (*D*(.,.)) within each cluster does not exceed a predefined threshold, *ϵ*. The last image of a cluster is the final image in the contiguous sequence for which the deformation does not exceed *ϵ*; including the next image exceeds this threshold. Each pair of successive clusters shares one image. Forward and backward transformations are computed within each cluster. The transformations enable estimation of deformation and identification of the consistently imaged region in the pre-hydrolysis frame, 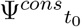, from which the consistently imaged cell walls, 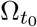, are derived. (B) The transformations are also used to propagate the pre-hydrolysis consistently imaged cell walls enabling the identification of consistently imaged cell walls in hydrolysis-phase image, denoted by 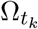, 0 *< k* ≤ *n*. (C) The transformations are then used to propagate the pre-hydrolysis segmentation, *S*_0_, to identify the cell walls in hydrolysis-phase images, 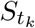, 0 *< k* ≤ *n*.

**Fig. S5:**
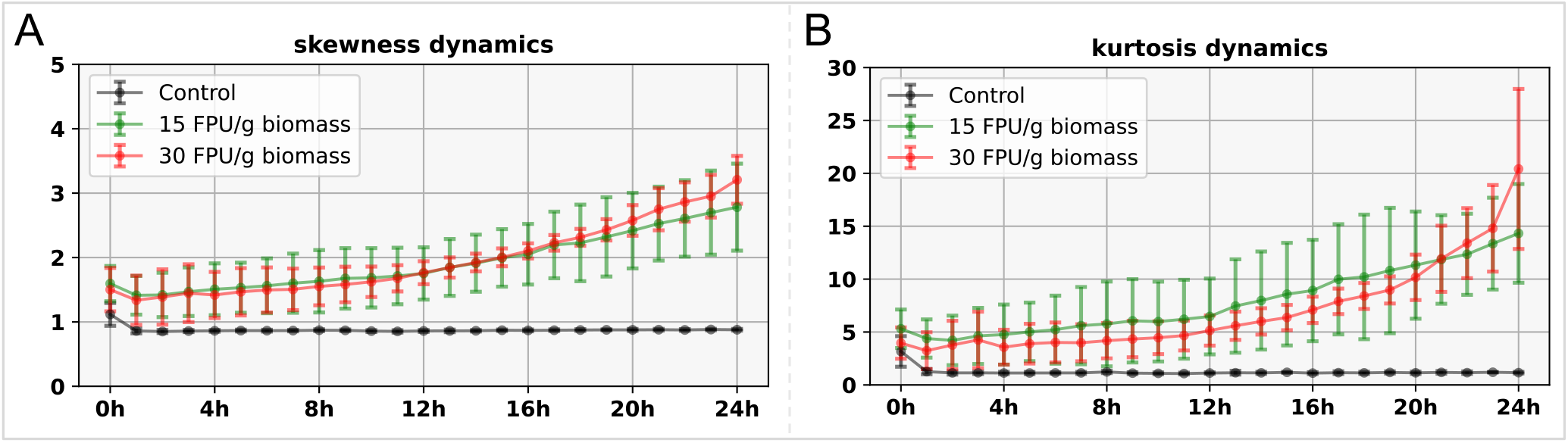
Skewness (A) and kurtosis(B) values of autofluorescence intensity distributions during enzymatic hydrolysis of control dataset and hydrolysis dataset collected with enzyme loadings of 15 and 30 FPU/g biomass.

## Notes

### Competing Interest Statement

The authors have declared no competing interest.

## References

[1] Abo, B.O., Gao, M., Wang, Y., Wu, C., Ma, H., Wang, Q., 2019. Lignocellulosic biomass for bioethanol: An overview on pretreatment, hydrolysis and fermentation processes. Rev. Environ. Health 34, 57–68. doi:10.1515/reveh-2018-0054.

[2] Ahamed, F., Song, H.S., Ooi, C.W., Ho, Y.K., 2019. Modelling heterogeneity in cellulose properties predicts the slowdown phenomenon during enzymatic hydrolysis. Chem. Eng. Sci. 206, 118–133. doi:10.1016/j.ces.2019.05.028.

[3] Auxenfans, T., Terryn, C., Päes, G., 2017. Seeing biomass recalcitrance through fluorescence. Sci. Rep. 7, 8838. doi:10.1038/s41598-017-08740-1.

[4] Bates, M., Huang, B., Dempsey, G.T., Zhuang, X., 2007. Multicolor Super-Resolution Imaging with Photo-Switchable Fluorescent Probes. Science 317, 1749–1753. doi:10.1126/science.1146598.

[5] Bhagia, S., Ďurkovič, J., Lagaňa, R., Kardosová, M., Kačík, F., Cernescu, A., Schäfer, P., Yoo, C.G., Ragauskas, A.J., 2022. Nanoscale FTIR and Mechanical Mapping of Plant Cell Walls for Understanding Biomass Deconstruction. ACS Sustain. Chem. Eng. 10, 3016–3026. doi:10.1021/acssuschemeng.1c08163.

[6] Binod, P., Janu, K., Sindhu, R., Pandey, A., 2011. Hydrolysis of Lignocellulosic Biomass for Bioethanol Production, in: Biofuels. Elsevier, pp. 229–250. doi:10.1016/B978-0-12-385099-7.00010-3.

[7] Bleuze, L., Lashermes, G., Alavoine, G., Recous, S., Chabbert, B., 2018. Tracking the dynamics of hemp dew retting under controlled environmental conditions. Ind. Crops. Prod. 123, 55–63. doi:10.1016/j.indcrop.2018.06.054.

[8] Blosse, S., Bouchoux, A., Montanier, C., Duru, P., 2023. X-ray microtomography reveals the 3D enzymatic deconstruction pathway of raw lignocellulosic biomass. Bioresour. Technol. Rep. 21, 101351. doi:10.1016/j.biteb.2023.101351.

[9] Cavelius, P., Engelhart-Straub, S., Mehlmer, N., Lercher, J., Awad, D., Brück, T., 2023. The potential of biofuels from first to fourth generation. PLOS Biol. 21, e3002063. doi:10.1371/journal.pbio.3002063.

[10] Chatterjee, C., Pong, F., Sen, A., 2015. Chemical conversion pathways for carbohydrates. Green Chem. 17, 40–71. doi:10.1039/C4GC01062K.

[11] Chen, Z., Chen, L., Khoo, K.S., Gupta, V.K., Sharma, M., Show, P.L., Yap, P.S., 2023. Exploitation of lignocellulosic-based biomass biorefinery: A critical review of renewable bioresource, sustainability and economic views. Biotechnol. Adv. 69, 108265. doi:10.1016/j.biotechadv.2023.108265.

[12] Cormen, T.H., Leiserson, C.E., Rivest, R.L., Stein, C., 2009. Introduction to Algorithms. MIT Press.

[13] Diao, Z., Ueda, K., Hou, L., Yamashita, H., Custance, O., Abe, M., 2023. Automatic drift compensation for nanoscale imaging using feature point matching. Appl. Phys. Lett. 122, 121601. doi:10.1063/5.0139330.

[14] Ding, D., Zhou, X., Ji, Z., You, T., Xu, F., 2016. How Does Hemicelluloses Removal Alter Plant Cell Wall Nanoscale Architecture and Correlate with Enzymatic Digestibility? Bioenergy Res. 9, 601–609. doi:10.1007/s12155-015-9703-1.

[15] Donaldson, L., 2020. Autofluorescence in Plants. Molecules 25, 2393. doi:10.3390/molecules25102393.

[16] Donaldson, L., Radotic, K., 2013. Fluorescence lifetime imaging of lignin autofluorescence in normal and compression wood. J. Microsc. 251, 178–187. doi:10.1111/jmi.12059.

[17] Dos Santos, A.C., Ximenes, E., Kim, Y., Ladisch, M.R., 2019. Lignin–Enzyme Interactions in the Hydrolysis of Lignocellulosic Biomass. Trends Biotechnol. 37, 518–531. doi:10.1016/j.tibtech.2018.10.010.

[18] Eibinger, M., Bubner, P., Ganner, T., Plank, H., Nidetzky, B., 2014. Surface structural dynamics of enzymatic cellulose degradation, revealed by combined kinetic and atomic force microscopy studies. FEBS J. 281, 275–290. doi:10.1111/febs.12594.

[19] Ghassemi, N., Poulhazan, A., Deligey, F., Mentink-Vigier, F., Marcotte, I., Wang, T., 2021. Solid-state NMR investigations of extracellular matrixes and cell walls of algae, bacteria, fungi, and plants. Chem. Rev. 122, 10036–10086. doi:10.1021/acs.chemrev.1c00669.

[20] Gierlinger, N., Keplinger, T., Harrington, M., 2012. Imaging of plant cell walls by confocal Raman microscopy. Nat. Protoc. 7, 1694–1708. doi:10.1038/nprot.2012.092.

[21] Guo, H., Zhao, Y., Chang, J.S., Lee, D.J., 2023. Enzymes and enzymatic mechanisms in enzymatic degradation of lignocellulosic biomass: A mini-review. Bioresour. Technol. 367, 128252. doi:10.1016/j.biortech.2022.128252.

[22] Gupta, R., Khasa, Y.P., Kuhad, R.C., 2011. Evaluation of pretreatment methods in improving the enzymatic saccharification of cellulosic materials. Carbohydr. Polym. 84, 1103–1109. doi:10.1016/j.carbpol.2010.12.074.

[23] Hallac, B.B., Ragauskas, A.J., 2011. Analyzing cellulose degree of polymerization and its relevancy to cellulosic ethanol. Biofuels Bioprod. Biorefining. 5, 215–225. doi:10.1002/bbb.269.

[24] Herbaut, M., Zoghlami, A., Habrant, A., Falourd, X., Foucat, L., Chabbert, B., Päes, G., 2018. Multimodal analysis of pretreated biomass species highlights generic markers of lignocellulose recalcitrance. Biotechnol. Biofuels. 11, 52. doi:10.1186/s13068-018-1053-8.

[25] Himmel, M.E., Ding, S.Y., Johnson, D.K., Adney, W.S., Nimlos, M.R., Brady, J.W., Foust, T.D., 2007. Biomass recalcitrance: engineering plants and enzymes for biofuels production. Science 315, 804–807. doi:10.1126/science.1137016.

[26] Hossein Khani, S., Ould Amer, K., Remy, N., Lebas, B., Habrant, A., Faraj, A., Malandain, G., Päes, G., Refahi, Y., 2025a. A distinct autofluorescence distribution pattern marks enzymatic deconstruction of plant cell wall. New Biotechnol. 88, 46–60. doi:10.1016/j.nbt.2025.04.001.

[27] Hossein Khani, S., Remy, N., Ould Amer, K., Lebas, B., Habrant, A., Malandain, G., Päes, G., Refahi, Y., 2025b. Time-lapse 3D image datasets of spruce tree wood enzymatic deconstruction. Data Brief 60, 111618. doi:10.1016/j.dib.2025.111618.

[28] Jang, S.K., Jeong, H., Choi, I.G., 2023. The Effect of Cellulose Crystalline Structure Modification on Glucose Production from Chemical-Composition-Controlled Biomass. Sustainability 15, 5869. doi:10.3390/su15075869.

[29] Kafle, K., Shin, H., Lee, C.M., Park, S., Kim, S.H., 2015. Progressive structural changes of Avicel, bleached softwood and bacterial cellulose during enzymatic hydrolysis. Sci. Rep. 5, 15102. doi:10.1038/srep15102.

[30] Kang, X., Kirui, A., Dickwella Widanage, M.C., Mentink-Vigier, F., Cosgrove, D.J., Wang, T., 2019. Lignin-polysaccharide interactions in plant secondary cell walls revealed by solid-state NMR. Nat. Commun. 10, 347. doi:10.1038/s41467-018-08252-0.

[31] Li, J., Kasal, B., 2025. Review on the Structure–Property Relationship of Lignocellulosic Materials Measured by Atomic Force Microscopy. Biomacromolecules 26, 1404–1418. doi:10.1021/acs.biomac.4c01278.

[32] Li, Y., Zhuo, J., Liu, P., Chen, P., Hu, H., Wang, Y., Zhou, S., Tu, Y., Peng, L., Wang, Y., 2018. Distinct wall polymer deconstruction for high biomass digestibility under chemical pretreatment in Miscanthus and rice. Carbohydr. Polym. 192, 273–281. doi:10.1016/j.carbpol.2018.03.013.

[33] Luterbacher, J.S., Moran-Mirabal, J.M., Burkholder, E.W., Walker, L.P., 2015. Modeling enzymatic hydrolysis of lignocellulosic substrates using fluorescent confocal microscopy II: Pretreated biomass. Biotechnol. Bioeng. 112, 32–42. doi:10.1002/bit.25328.

[34] Ma, H., Chen, M., Nguyen, P., Liu, Y., 2024. Toward drift-free high-throughput nanoscopy through adaptive intersection maximization. Sci. Adv. 10, eadm7765. doi:10.1126/sciadv.adm7765.

[35] Maceda, A., Terrazas, T., 2022. Fluorescence Microscopy Methods for the Analysis and Characterization of Lignin. Polymers 14, 961. doi:10.3390/polym14050961.

[36] Malik, K., Sharma, P., Yang, Y., Zhang, P., Zhang, L., Xing, X., Yue, J., Song, Z., Nan, L., Yujun, S., El-Dalatony, M.M., Salama, E.S., Li, X., 2022. Lignocellulosic biomass for bioethanol: Insight into the advanced pretreatment and fermentation approaches. Ind. Crops. Prod. 188, 115569. doi:10.1016/j.indcrop.2022.115569.

[37] McCann, M.C., Carpita, N.C., 2015. Biomass recalcitrance: A multi-scale, multi-factor, and conversion-specific property. J. Exp. Bot. 66, 4109–4118. doi:10.1093/jxb/erv267.

[38] Meng, X., Foston, M., Leisen, J., DeMartini, J., Wyman, C.E., Ragauskas, A.J., 2013. Determination of porosity of lignocellulosic biomass before and after pretreatment by using Simons’ stain and NMR techniques. Bioresour. Technol. 144, 467–476. doi:10.1016/j.biortech.2013.06.091.

[39] Meng, X., Wells, T., Sun, Q., Huang, F., Ragauskas, A., 2015. Insights into the effect of dilute acid, hot water or alkaline pretreatment on the cellulose accessible surface area and the overall porosity of Populus. Green Chem. 17, 4239–4246. doi:10.1039/C5GC00689A.

[40] Michelin, G., Refahi, Y., Wightman, R., Jonsson, H., Traas, J., Godin, C., Malandain, G., 2016. Spatio-temporal registration of 3D microscopy image sequences of arabidopsis floral meristems, in: 2016 IEEE 13th International Symposium on Biomedical Imaging (ISBI), IEEE, Prague, Czech Republic. pp. 1127–1130. doi:10.1109/ISBI.2016.7493464.

[41] Moreira-Vilar, F.C., Siqueira-Soares, R.d.C., Finger-Teixeira, A., Oliveira, D.M.d., Ferro, A.P., da Rocha, G.J., Ferrarese, M.d.L.L., dos Santos, W.D., Ferrarese-Filho, O., 2014. The acetyl bromide method is faster, simpler and presents best recovery of lignin in different herbaceous tissues than klason and thioglycolic acid methods. PLoS One 9, e110000. doi:10.1371/journal.pone.0110000.

[42] Muzamal, M., Jedvert, K., Theliander, H., Rasmuson, A., 2015. Structural changes in spruce wood during different steps of steam explosion pretreatment. Holzforschung 69, 61–66. doi:10.1515/hf-2013-0234.

[43] Netherer, S., Lehmanski, L., Bachlehner, A., Rosner, S., Savi, T., Schmidt, A., Huang, J., Paiva, M.R., Mateus, E., Hartmann, H., Gershenzon, J., 2024. Drought increases Norway spruce susceptibility to the Eurasian spruce bark beetle and its associated fungi. New Phytol. 242, 1000–1017. doi:10.1111/nph.19635.

[44] Ourselin, S., Roche, A., Prima, S., Ayache, N., 2000. Block matching: A general framework to improve robustness of rigid registration of medical images, in: International Conference on Medical Image Computing And Computer-Assisted Intervention, Springer. pp. 557–566. doi:10.1007/978-3-540-40899-4_57.

[45] Peng, P., She, D., 2014. Isolation, structural characterization, and potential applications of hemicelluloses from bamboo: A review. Carbohydr. Polym. 112, 701–720. doi:10.1016/j.carbpol.2014.06.068.

[46] Pérez, J., Muñoz-Dorado, J., De La Rubia, T., Martínez, J., 2002. Biodegradation and biological treatments of cellulose, hemicellulose and lignin: An overview. Int. Microbiol. 5, 53–63. doi:10.1007/s10123-002-0062-3.

[47] Qaseem, M.F., Shaheen, H., Wu, A.M., 2021. Cell wall hemicellulose for sustainable industrial utilization. Renew. Sustain. Energy Rev. 144, 110996. doi:10.1016/j.rser.2021.110996.

[48] Refahi, Y., Zardilis, A., Michelin, G., Wightman, R., Leggio, B., Legrand, J., Faure, E., Vachez, L., Armezzani, A., Risson, A.E., et al., 2021. A multiscale analysis of early flower development in arabidopsis provides an integrated view of molecular regulation and growth control. Dev. Cell 56, 540–556. doi:10.1016/j.devcel.2021.01.019.

[49] Refahi, Y., Zoghlami, A., Viné, T., Terryn, C., Päes, G., 2024. Plant cell wall enzymatic deconstruction: Bridging the gap between micro and nano scales. Bioresour. Technol. 414, 131551. doi:10.1016/j.biortech.2024.131551.

[50] Shi, X., Wang, L., Sun, L., Chen, H., 2022. Sufficient premixing enhances enzymatic hydrolysis efficiency of lignocellulose at high-solids loading. Chem. Eng. J. 444, 136612. doi:10.1016/j.cej.2022.136612.

[51] Simmons, T.J., Mortimer, J.C., Bernardinelli, O.D., Pöppler, A.C., Brown, S.P., deAzevedo, E.R., Dupree, R., Dupree, P., 2016. Folding of xylan onto cellulose fibrils in plant cell walls revealed by solid-state NMR. Nat. Commun. 7, 13902. doi:10.1038/ncomms13902.

[52] Singh, S.S., Lim, L.T., Manickavasagan, A., 2023. Imaging and spectroscopic techniques for microstructural and compositional analysis of lignocellulosic materials: A review. Biomass Convers. Bioref. 13, 499–517. doi:10.1007/s13399-020-01075-4.

[53] Velvizhi, G., Balakumar, K., Shetti, N.P., Ahmad, E., Kishore Pant, K., Aminabhavi, T.M., 2022. Integrated biorefinery processes for conversion of lignocellulosic biomass to value added materials: Paving a path towards circular economy. Bioresour. Technol. 343, 126151. doi:10.1016/j.biortech.2021.126151.

[54] Vicente, N.B., Zamboni, J.E.D., Adur, J.F., Paravani, E.V., Casco, V.H., 2007. Photobleaching correction in fluorescence microscopy images. J. Phys.: Conf. Ser. 90, 012068. doi:10.1088/1742-6596/90/1/012068.

[55] VTK, The Visualization Toolkit. [oneline]. Available: https://vtk.org.

[56] Yang, B., Dai, Z., Ding, S.Y., Wyman, C.E., 2011. Enzymatic hydrolysis of cellulosic biomass. Biofuels 2, 421–449. doi:10.4155/bfs.11.116.

[57] Yoo, C.G., Meng, X., Pu, Y., Ragauskas, A.J., 2020. The critical role of lignin in lignocellulosic biomass conversion and recent pretreatment strategies: A comprehensive review. Bioresour. Technol. 301, 122784. doi:10.1016/j.biortech.2020.122784.

[58] Zeng, Y., Zhao, S., Yang, S., Ding, S.Y., 2014. Lignin plays a negative role in the biochemical process for producing lignocellulosic biofuels. Curr. Opin. Biotechnol. 27, 38–45. doi:10.1016/j.copbio.2013.09.008.

[59] Zhao, W., Fernando, L.D., Kirui, A., Deligey, F., Wang, T., 2020. Solid-state NMR of plant and fungal cell walls: A critical review. Solid State Nucl. Magn. Reson. 107, 101660. doi:10.1016/j.ssnmr.2020.101660.

[60] Zoghlami, A., Refahi, Y., Terryn, C., Päes, G., 2019. Multimodal characterization of acid-pretreated poplar reveals spectral and structural parameters strongly correlate with saccharification. Bioresour. Technol. 293, 122015. doi:10.1016/j.biortech.2019.122015.

[61] Zoghlami, A., Refahi, Y., Terryn, C., Päes, G., 2020. Three-Dimensional Imaging of Plant Cell Wall Deconstruction Using Fluorescence Confocal Microscopy. Sustain. Chem. 1, 75–85. doi:10.3390/suschem1020007.

